# Numb reduces Tau levels and prevents neurodegeneration in mouse models of tauopathy in an isoform-specific manner

**DOI:** 10.1101/2021.09.28.462203

**Authors:** Marine Lacomme, Sarah C. Hales, Katarina Stevanovic, Christine Jolicoeur, Therence Bois, Jenny Cai, Melissa Desrosiers, Deniz Dalkara, Michel Cayouette

## Abstract

Accumulation of the microtubule-associated protein Tau is linked to neuronal cell death in tauopathies, but how exactly intraneuronal Tau levels are regulated in health and disease remains unclear. Here we identify the trafficking adaptor protein Numb as an essential regulator of Tau homeostasis. Conditional inactivation of Numb in retinal ganglion cells (RGCs) increases monomeric and oligomeric Tau levels, leading to axonal blebbing followed by neuronal cell loss in aged mice. Moreover, in a mouse model of tauopathy, inactivation of Numb in RGCs and spinal motoneurons accelerates neurodegeneration, and leads to precocious hindlimb paralysis. Conversely, overexpression of the long isoform of Numb (Numb-72), but not other isoforms, decreases intracellular Tau levels by promoting the extracellular release of monomeric Tau, and AAV-mediated delivery of Numb-72 in RGCs *in vivo* prevents neurodegeneration in two different mouse models of tauopathy. Taken together, these results uncover Numb as a modulator of intracellular Tau levels and identify Numb-72 as a novel therapeutic factor for tauopathies.

## INTRODUCTION

The microtubule binding protein Tau is a soluble protein that stabilizes axonal microtubules in healthy neurons. In a group of neurodegenerative diseases called tauopathies, however, post-translational modifications generate insoluble Tau, which can lead to the formation of toxic oligomers and neurofibrillary tangles (*1–3*). In recent years, accumulation of Tau oligomers has been shown to contribute directly to neurodegeneration (*4–7*), leading to the idea that lowering intracellular levels of Tau before it forms oligomers might alleviate the toxic effects of oligomers and prevent neuronal cell death in tauopathies. But would decreasing the levels of native Tau proteins produce undesirable side effects? Surprisingly, previous experiments have shown that inactivation of *Mapt*, the gene coding for Tau, is relatively well tolerated in mice (*8*) (*9*) (*10*), suggesting that Tau-lowering approaches may be viable. Accordingly, reducing Tau protein levels in various ways, such as antisense oligonucleotides, stimulation of proteasomal degradation and autophagy, or manipulation of a kinase regulating Tau stability was found to prevent or even reverse neurological phenotypes in tauopathy human neurons and mouse models (*11–14*) (*15*). While these studies demonstrate the therapeutic potential of Tau-lowering approaches, how exactly Tau levels are normally regulated at steady state remains largely unclear. Therefore, we aimed to identify natural regulators of Tau trafficking, which we postulated would not only advance our understanding of how Tau levels are regulated in healthy neurons and how it might become dysregulated in disease, but also provide novel ways to lower Tau levels and protect neurons in tauopathy.

Regulation of intracellular trafficking plays a critical part in protein homeostasis. One adaptor protein that was found to regulate intracellular trafficking in the nervous system is Numb. In Drosophila sensory organ precursors, d-numb regulates cell fate decisions by decreasing Notch signaling (*16*). In vertebrates, Numb and Numb-like (Nbl) are homologs of d-numb and display extensive sequence homology and functional redundancy, including inhibition of Notch signaling via ubiquitination of the Notch receptor and trafficking of the intracellular domain for proteasome-mediated degradation (*17, 18*). Numb/Nbl were also found to play an important role in cell fate decisions during neural progenitor asymmetric divisions (*19–21*), neuronal migration (*22*), neurite outgrowth (*23, 24*) and axon guidance (*25*). Classically referred to as an antagonist of Notch signaling, Numb has since been involved in the trafficking of many different cargo proteins (*26, 27*). Alternative splicing of Numb leads to the production of four different isoforms in mice, defined by their short or long phosphotyrosine-binding domain (PTB) and proline-rich region (PRR), which are thought to mediate different functions. Interestingly, a previous study showed that Numb isoforms with a short PTB (Numb-65 and Numb-71) are elevated in the parietal cortex of Alzheimer’s patients (*28*), suggesting that a correct balance between the long (Numb-72 and Numb-66) and short PTB isoforms may play a role in neuronal survival and degeneration. However, while Numb function has been studied in various neurodevelopmental contexts, its role in mature neurons remains largely unexplored. Given the widespread expression of Numb in the adult CNS and its role in intracellular trafficking, we hypothesized that it may be involved in regulating Tau homeostasis, thereby promoting neuronal survival.

To test this hypothesis, we primarily used the mouse retina as a model system. In Alzheimer’s disease and other tauopathies, retinal ganglion cells (RGCs), the projection neurons of the retina, display elevated Tau levels and eventually degenerate (*29–33*). Consistently, Tau overexpression in mouse RGCs triggers cell death (*34, 35*), and mouse models of tauopathy display RGC degeneration (*33, 36*). Therefore, similar to brain neurons, RGCs are highly sensitive to an increase in Tau levels. Combined with the accessibility and anatomical simplicity of the mouse retina, this makes RGCs particularly attractive to study the role of Numb in the regulation of Tau in CNS neurons. Using conditional mouse genetics and tauopathy disease models, we report here that Numb is an essential regulator of intraneuronal Tau levels and prevents degeneration of tauopathy neurons in an isoform-specific manner.

## RESULTS

### Numb is required for long-term neuronal survival

We previously reported that *Numb* and *Nbl* transcripts are both expressed in the RGC layer of the adult mouse retina (*37*). As the RGC layer also contains displaced amacrine cells and astrocytes, we first sought to determine more precisely whether Numb and Nbl proteins are actually expressed in RGCs by co-staining adult retinal sections for Brn3b, a pan-RGC marker, together with an antibody that recognizes both Numb and Nbl (*38*) (Fig. S1 A-C). As expected, Numb/Nbl were detected in RGCs, including their axons and dendrites, although the strongest signal was observed in the soma (Fig. S1 A-E).

To uncover a potential role for Numb/Nbl in RGCs, we crossed the Islet1-Cre driver mouse line (*39*) to the Numb floxed line, in which exon 1 of the *Numb* gene is flanked by loxP sites (*40*). As *Nbl* is known to compensate for the loss of *Numb* in multiple contexts (*41, 42*), we generated the *Numb* conditional knockout mouse on a *Nbl* null background (*43*), allowing us to study the specific role of *Numb* in RGCs (Fig. S1 F). To assess Cre activity, we additionally introduced a Rosa26-loxP-STOP-loxP-tdTomato (tdT) reporter allele. As expected, tdT expression was observed in Islet1-expressing cells in the retina, such as RGCs, from as early as embryonic day 14.5, as well as bipolar cells and a subset of amacrine cells (*44*) (Fig. S1 G-L) In this study, Islet1^Cre/+^; Numb^fl/fl^; Numbl^del/del^ animals will be referred to as conditional double knockouts (cDKO), Islet1^Cre/+^; Numb^fl/+^; Numbl^del/del^ will be referred to as Numb^+/−^;Nbl KO and Islet1^+/+^; Numb^fl/fl^; Numbl^del/del^ will be referred as Nbl KO. As expected, we found a loss of Numb immunostaining signal in cDKO RGCs compared to Numb^+/−^; Nbl KO (Fig. S1 M-P). As Nbl is null in both conditions, we conclude that the immunostaining signal observed in RGCs is specific for Numb. We also performed PCR on genomic DNA of adult retinas from wild-type (WT), Numb^+/−^; Nbl KO, Nbl KO and cDKO (Fig. S1 Q-T). The excision was not complete, but this was expected since a large proportion of cells expressing Numb in the retina do not carry the Islet-Cre driver. These results confirm the generation of a mouse line that allows Cre-mediated inactivation of Numb in RGCs.

To study the role of Numb on neuronal survival, we first analyzed retinal sections from 5- and 20-month-old cDKO compared to Numb^+/−^;Nbl KO and Nbl KO controls (Fig. S2 A-K). At 5 months, stainings for the RGC marker Brn3b, the amacrine cell marker Pax6, or the bipolar cell marker Chx10 revealed no significant changes in the number of these cell types in cDKO retinas compared to controls (Fig. S2 I-K). In 20-month-old mice (Fig. S2 L-V), however, we observed a 3-fold reduction in the number of RGCs in cDKOs (Fig. S2 V), whereas the number of amacrine and bipolar cells was unchanged (Fig. S2 T and U). As *Nbl* is null in all conditions examined, these results indicate that Numb is specifically required for the long-term survival of RGCs, but dispensable for bipolar and amacrine cell survival. To reinforce this finding and determine the time-course of RGC loss, we prepared retinal flatmounts from 5-, 8- and 20-month old mice and counted the number of Brn3b+ RGCs (Fig. 1 M). At 5 months, the total number of RGCs was similar in all genotypes examined and not significantly different from WT mice, consistent with our observations in retinal sections (Fig.1 A-D, M and Fig. S2K). Starting in 8-month-old animals, however, we observed a significant decrease in RGC numbers in cDKO animals (Fig.1 E-H, M), which progressed to about 50% loss of RGCs at 20 months (Fig.1 I-L, M). Importantly, the number of RGCs in Nbl KO mice was not different from that of WT animals at any age examined, indicating that Nbl inactivation alone has no effect on RGC survival (Fig. 1M). Taken together, these results show that Numb is required to support the long-term survival of RGCs.

**Fig. 1.**
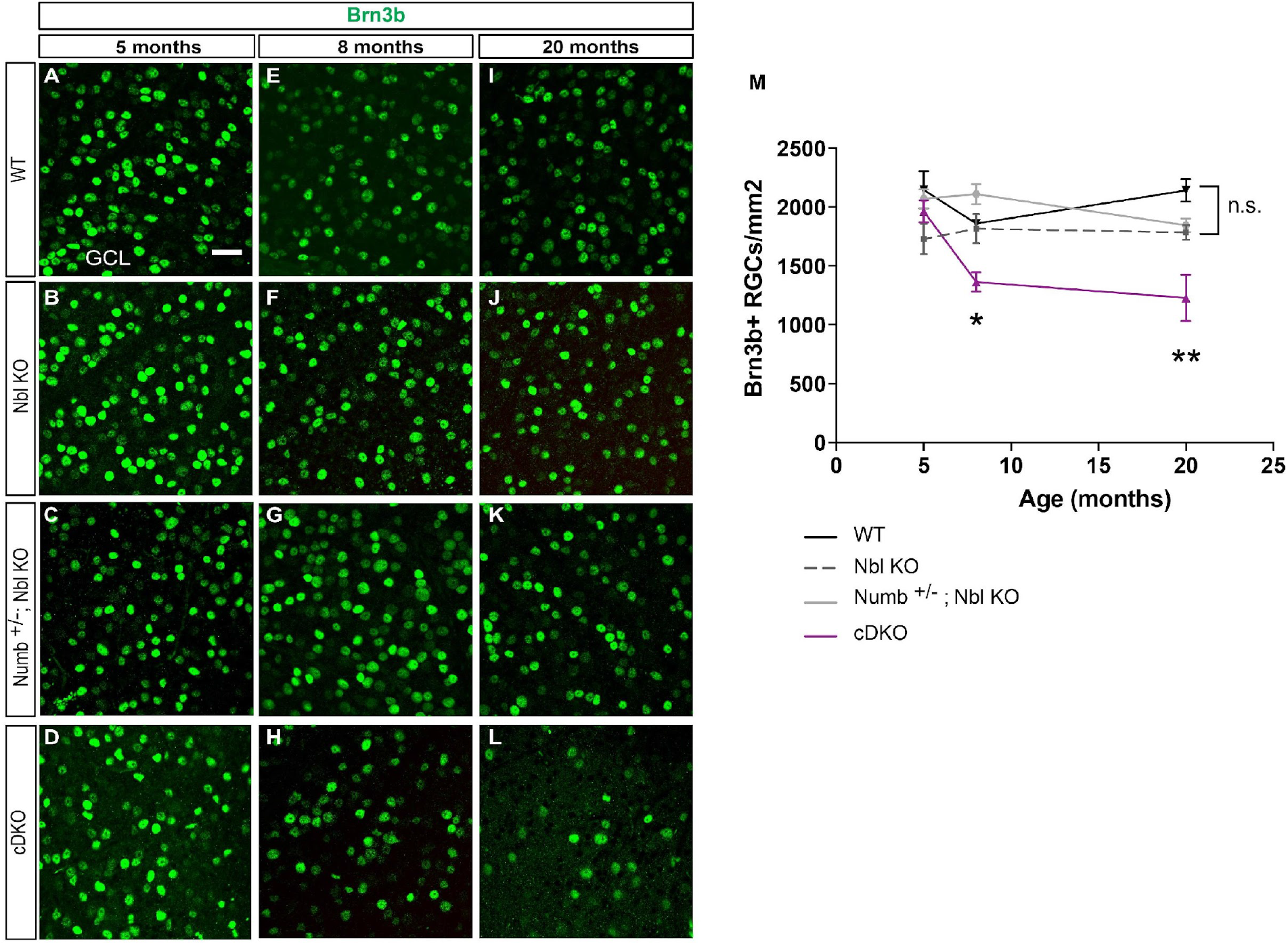
Numb is essential for long-term survival of RGCs. **(A-L)** Single-plane confocal images of retinal flat mounts stained for Brn3b from 5-month-old (A-D), 8-months-old (E-H) and 20-month-old (I-L) WT, Nbl KO, Numb^−/+^; Nbl KO and cDKO mice, as indicated. Images were taken in the ganglion cell layer (GCL). Scale bar = 30μm **(M)** Quantification of the number of Brn3b+ RGC per mm^2^ in WT, controls and cDKO mice at 5, 8 and 20 months old. Mean ± SEM, n= 5 animals/genotype/time point. * p≤0.05, **p≤0.01, two-way Anova followed by Turkey’s test. n.s: not significant.

### Numb is required to maintain axonal integrity

In many neurodegenerative diseases, damage occurs in axons and dendrites before any cell death is detected (*45, 46*). To test whether this was the case in cDKO retinas, we evaluated the morphology of RGC axons in 5-month-old animals, prior to any detectable cell loss (Fig. 1 M). First, we used DiI to retrogradely label RGC axons in Nbl KO and cDKO mice. While axons of Nbl KO animals appeared smooth and healthy, those from cDKO mice showed extensive blebbing (Fig. 2A–D), an early sign of neurodegeneration. Second, we used adeno-associated viral vector type 2 expressing GFP (AAV2-GFP), which preferentially infects RGCs after intravitreal injections(*47*), to label RGC axons in Nbl KO or cDKO animals. Similar to what we observed with DiI, axonal blebbing was extensive in cDKO RGCs, but not in Nbl KO (Fig. 2E, F). We next asked whether these axonal defects were cell autonomous. To do this, we cultured primary RGCs from postnatal day 8 (P8) Numb^+/−^; Nbl KO and cDKO mice and counted the number of axonal blebs 14 days later using the tdT reporter and immunostaining for neurofilament 165kDa subunit (NF165), a marker of RGC axons. As observed *in vivo*, RGCs from cDKO mice had significantly more blebs compared to Numb^+/−^; Nbl KO, whereas total neurite length and number of branches was not affected (Fig. 2 G–N). Together, these results indicate that axonal blebbing occurs prior to RGC degeneration in cDKO mice, and that Numb functions cell-autonomously to maintain axonal integrity in RGCs.

**Fig. 2.**
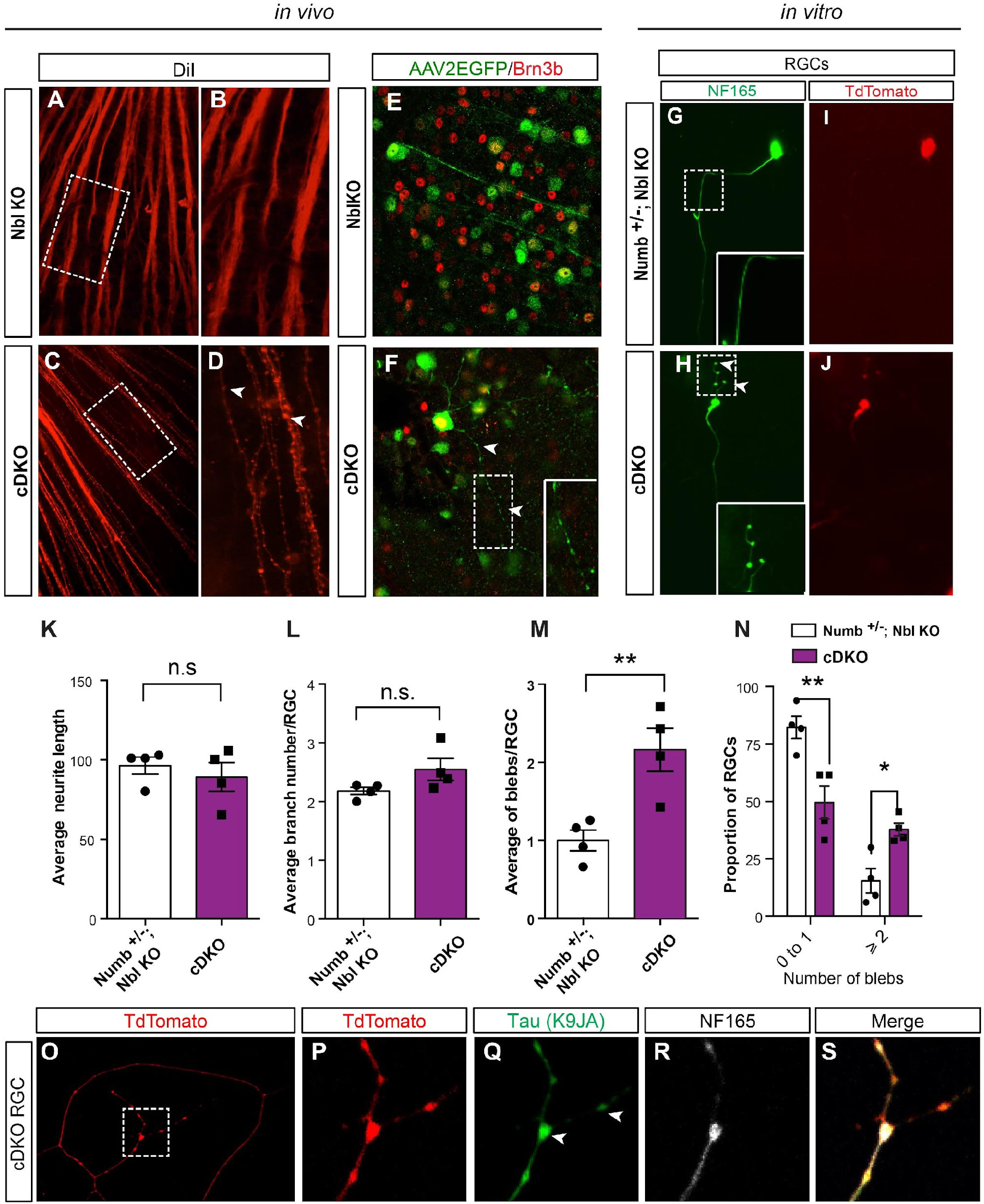
Numb is essential to maintain axonal homeostasis. **(A-D)** Confocal images of DiI-labelled RGC axons in retinal flatmounts from 5-month-old Nbl KO (A, B) and cDKO (C, D). Higher magnification views are shown in B and D. Arrowheads point to blebbing. Images in (B) and (D) show high magnification views of the regions identified by the dotted box in (A) and (C). **(E, F)** Confocal images of AAV2EGFP-infected Nbl KO (E) and cDKO (F) RGCs immunostained for Brn3b (red) on retinal flatmounts. Arrowheads point to axonal blebs. Magnified view of the boxed region are shown in the inset in (G) and (H). **(G-N)** Primary retinal cell cultures prepared from Numb^+/−^; Nbl KO and cDKO immunostained for neurofilament-165 (NF165) and TdTomato at 14 days in culture. Blebbing (arrowheads) is observed in cDKO RGCs. Insets show magnified views of the boxed area. **(K-N)** Quantification of neurite length (K), branch number (L) and number of axonal blebs (M-N) in Numb^+/−^; Nbl KO and cDKO RGCs. Number of blebs in controls are normalized to 1 in (M). Graphs show mean ± SEM, *p<0.05; ** p ≤0.001; n.s. non-significant, Student’s t test. n=4 independent cultures. A total of 122 neurons were counted in Numb^−/+^; Nbl KO and 155 neurons in cDKO. Graph in (N) shows repartition of the number of blebs in different conditions. Two-way Anova followed by Turkey’s test, mean ± SEM, *p<0.05; ** p ≤0.001; n.s. not significant. n=4 independent cultures. A total of 122 neurons were counted in Numb^−/+^; Nbl KO and 155 neurons in cDKO. **(O-S)** Immunostaining for total Tau (K9JA), NF165, and TdTomato on primary retina cell cultures prepared from cDKO retinas. Images in P-S represent the zoomed-in view of the boxed area in O. Axonal blebs contain Tau (arrowhead).

### Numb negatively controls intraneuronal Tau levels

In tauopathies, the accumulation of Tau proteins is often associated with the formation of axonal blebs (*48, 49*), similar to what we observed in cDKO RGCs. Consistently, we found that the axonal blebs observed in cDKO RGCs are filled with Tau (Fig. 2Q). We therefore hypothesized that cDKO RGCs might have abnormal levels of Tau. To test this idea, we prepared optic nerve protein extracts from 5-month-old cDKO animals and assessed Tau levels by Western blot. We found that the levels of total Tau in cDKO optic nerves is more than 6-fold higher than in Nbl KO optic nerves (Fig. 3A, B), whereas acetylated tubulin was not changed (Fig. 3A), suggesting that the elevation in Tau levels is not a consequence of destabilized microtubules. Importantly, the levels of toxic Tau oligomers were also sharply increased in cDKO optic nerves (Fig. 3C–E). Of note, we found no changes in the overall levels of phosphorylated Tau (p-Tau) in cDKO optic nerve (Fig. S3 A, B) and retina (Fig. S3 C, D) extracts compared to Nbl KO, suggesting that elevation of Tau oligomers in cDKO occurs in a phosphorylation-independent manner. Moreover, we did not find any modification in Tau mRNA levels in cDKO (Fig. S3 E), indicating that Numb does not regulate Tau transcription. Altogether, these results show that Numb is required to negatively regulate total Tau protein levels in RGCs and prevent the accumulation of oligomeric Tau.

**Fig. 3.**
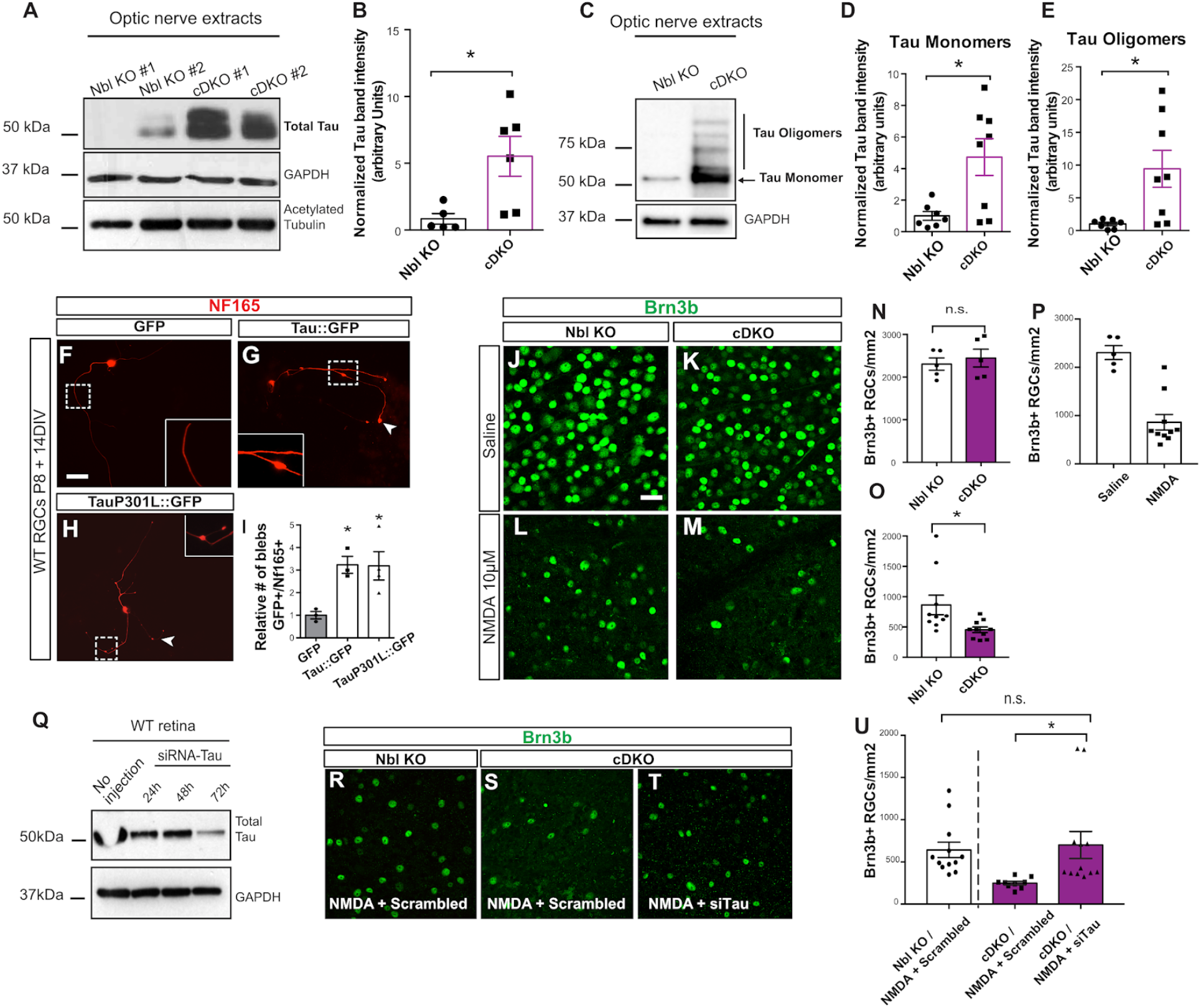
Tau levels are elevated in Numb cDKO optic nerves and cause RGC death. **(A)** Western blot analysis of total Tau (5A6), GAPDH and acetylated tubulin expression in optic nerve extracts prepared from 5-month-old Nbl KO and cDKO mice. **(B)** Quantification of the levels of Tau in western blots, relative to GAPDH. Graph shows mean ± SEM, * p ≤0.05; Student’s t test, n=7 Nbl KO and n=7 cDKO. **(C)** Western blot analysis of Tau monomer (50kDa) and oligomer (>50 kDa) expression using the T22 antibody on optic nerve extracts prepared from 5-month-old Nbl KO and cDKO mice. **(D-E)** Quantification of relative levels of monomers (D) and oligomers (E) of Tau from western blots. GAPDH expression was used as loading control. Graph shows mean ± SEM, *p ≤0.05; Student’s t test, n=7 Nbl KO and 8 cDKO. **(F-H)** Primary retinal cell cultures 14 days after transfection with GFP (F), Tau∷GFP (G) and TauP301L∷GFP (H) stained for NF165 (red). Arrowheads point to blebs. **(I)** Quantification of the number of blebs/RGC in the different conditions, as indicated. Graph shows mean ± SEM, * p ≤0.05; Anova one-way test followed by Dunnett’s test. n=3 independent cultures. Total number of cells counted: 101 for GFP, 92 for Tau∷GFP, and 82 for TauP301L∷GFP. **(J-M)** Immunostaining for Brn3b on retinal flatmounts from 5-month-old Nbl KO and cDKO mice 3 days after injection of saline (J, K) or NMDA (10mM; L,M). Images were taken in the ganglion cell layer. Scale bar= 30μm. **(N, O)** Quantification of the number of Brn3b+ RGCs per mm^2^ in Nbl KO and cDKO mice 3 days after intravitreal injection of saline (N) or NMDA (O). Graph shows mean ± SEM, * p ≤0.05, n.s. not significant, Student’s t test. n=5 animals/genotype with saline and n=10animals/genotype with NMDA **(P)** Quantification of the number of Brn3b+ RGCs per mm^2^ in WT mice 3 days after intravitreal injection of saline or NMDA. Graph shows mean ± SEM. n=5 animals with saline injection and n=10animals with NMDA injection. **(Q)** Western blot analysis of total Tau and GAPDH expression in retinal extracts prepared 24h, 48h or 72h after intravitreal injection of siRNA against Tau. **(R-T)** Immunostaining for Brn3b on retinal flat mounts from 5-months-old Nbl KO and cDKO mice 72h after intravitreal injection of NMDA (10mM) and scramble siRNA (S) or with siTAU (T). Confocal images were taken in the ganglion cell layer. Scale bar = 30μm. **(U)** Quantification of number of Brn3b RGCs per mm^2^ in Nbl KO and cDKO mice 72h after injection of NMDA and siRNA. Graph shows mean ± SEM, n.s: not significant, * p ≤0.05; Anova one-way test followed by Turkey’s test. n=12 NblKO, n=9 cDKO NMDA + Scramble and n=12 cDKO NMDA + siTau.

Based on the above results, we postulated that the increased Tau levels observed in RGC axons might be the cause of neurodegeneration in cDKO retinas. If this were the case, we predicted that elevating Tau levels in WT RGCs would phenocopy the loss of Numb, whereas knocking down Tau in cDKO RGCs would rescue degeneration. To test these predictions, we first overexpressed GFP-fused WT human Tau (Tau∷GFP) or TauP301L (TauP301L∷GFP), a mutant form of Tau associated with tauopathies, in cultured WT mouse RGCs and quantified axonal blebs 14 days later. As predicted, overexpression of Tau∷GFP or TauP301L∷GFP significantly increased axonal blebbing compared to GFP expression alone (Fig. 3F–I), indicating that the overall levels of Tau are critical to RGC axonal integrity. Next, we wanted to determine whether lowering Tau levels could rescue degeneration of cDKO RGCs *in vivo.* As shown in Figure 1, however, significant loss of neurons is not detected before 8 months of age in cDKO mice, rendering this experiment difficult. To circumvent this issue, we sought to accelerate cell death in cDKO animals. To do this, we injected low doses of NMDA in the eyes of Nbl KO and cDKO mice to induce a mild excitotoxic stress (*50–52*). At this dose of NMDA, the number of Brn3b+ RGCs was reduced by 50-60% in Nbl KO three days after injection, compared to saline-injected controls (Fig. 3J–O), whereas the same dose reduced RGC numbers by an additional 50% in cDKO mice (Fig. 3 M, O). Thus, cDKO RGCs are more susceptible to stress-induced degeneration than controls, providing a fast and convenient way to study Numb-dependent neurodegeneration. Using this assay, we injected a validated Tau siRNA cocktail (Fig. 3Q) or scrambled siRNA together with NMDA into the vitreous of Nbl KO or cDKO mice and compared RGC numbers three days later. We found that knocking-down Tau completely rescues neurodegeneration in cDKO RGCs (Fig. 3 R–U). These results indicate that elevated Tau levels are responsible for neurodegeneration in cDKO mice.

### Numb delays Tau-mediated neurodegeneration in a mouse model of tauopathy

The above results indicate that Numb is required to control intracellular levels of Tau and promote neuronal survival. In light of these findings, we postulated that loss of Numb might exacerbate neuronal degeneration in a model of tauopathy, wherein elevated Tau levels correlate with neuronal cell loss (*53, 54*). To test this hypothesis, we generated a mouse line in which *Numb* and *Nbl* are inactivated on a TauP301S transgenic background (cDKO; TauP301S), a mouse model of tauopathy in which Tau levels in RGCs are five times those observed in control mice(*7*). We then compared RGC numbers at 8 months-old in Nbl KO; TauWT, which have the same number of RGCs as WT mice (see Fig. 1M), to that of i) Nbl KO; TauP301S, ii) cDKO; TauWT, and iii) cDKO; TauP301S. While Nbl KO; TauP301S and cDKO; TauWT mice had a similar reduction of RGC numbers, we found that cDKO; TauP301S mice have an even greater reduction of RGCs (Fig. 4 E). These results show that Numb functions to slow down RGC degeneration in the context of tauopathies.

**Fig. 4.**
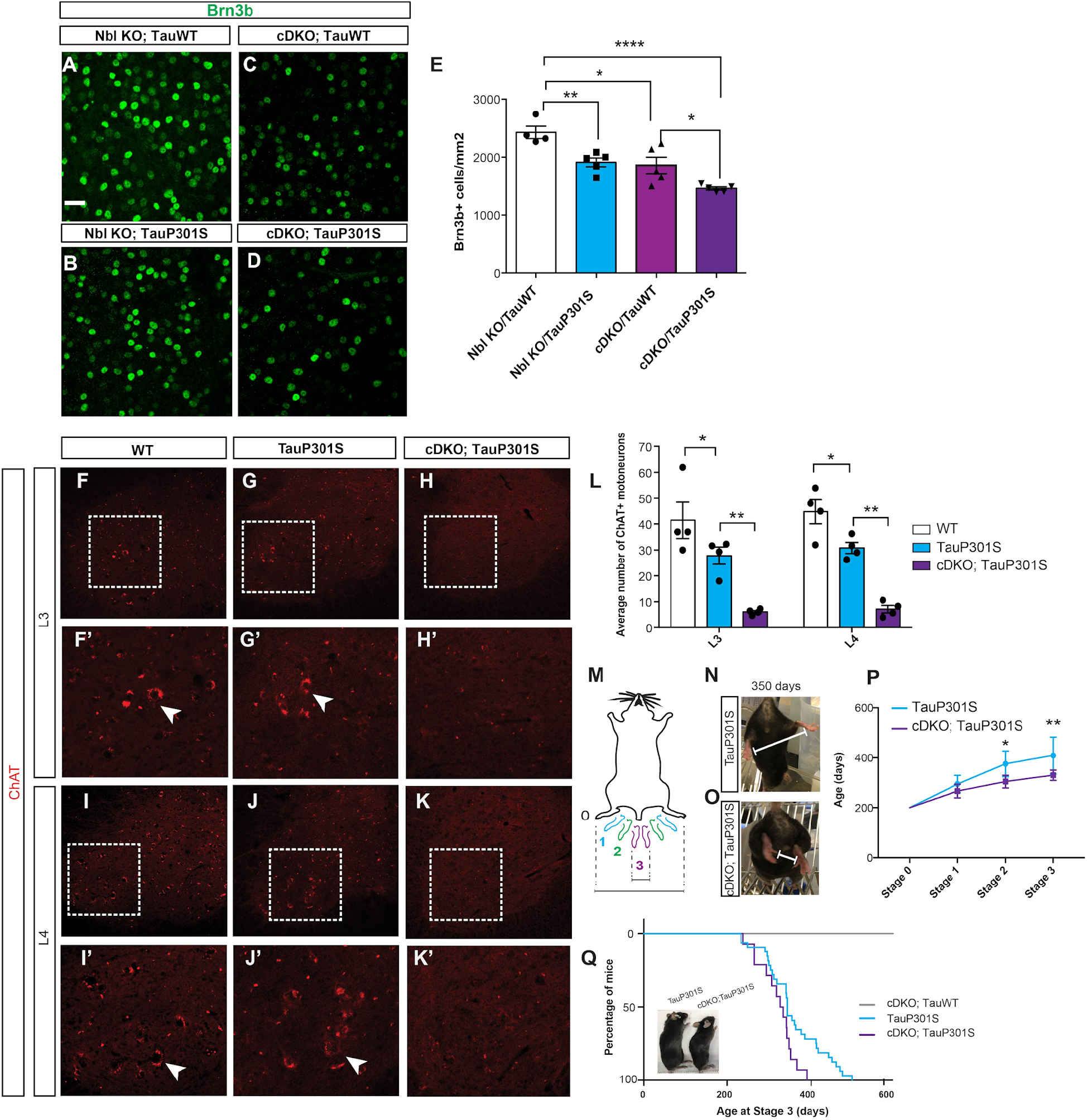
Loss of Numb in TauP301S mouse accelerates neuronal degeneration. **(A-D)** Immunostaining for Brn3b on retina flat mounts from 8-month-old Nbl KO; TauWT (A), Nbl KO; TauP301S (B), cDKO; TauWT (C), and cDKO;TauP301S (D). Scale bar = 30μm. **(E)** Quantification of the number of Brn3b+ RGCs per mm^2^ in 8-months-old cDKO;TauP301S and the various controls, as indicated. Graph shows mean ± SEM, n= 5 animals/genotype. One-way Anova followed by Turkey’s test, * p≤0.05, **p≤0.01, ****p≤0.0001. **(F-K)** Immunostaining for choline acetyltransferease (ChAT) on adult spinal cord sections in the lumbar region L3 (F-H) and L4 (I-K) of WT (F, F’ and I,I’), TauP301S (G, G’ and J,J’) and cDKO; TauP301S (H, H’ and K,K’) mice at 350 days. Arrows point to ChAT+ cells. (**L)** Quantification of the average number of ChAT+ motoneurons in L3 and L4 in WT, TauP301S and cDKO; TauP301S. Graph shows mean ± SEM, n= 4 animals/genotype. Two-way Anova followed by Turkey’s test, * p≤0.05, **p≤0.01. **(M)** Schematic representation of paralysis stages based on the clasping reflex, 0 corresponds to no paralysis and 3 to full paralysis of the hindlimbs. **(N-O)** Representative image of 350-day-old transgenic TauP301S mouse showing no paralysis (stage 0) and normal hindlimb clasping reflex (N) or cDKO; TauP301S mouse with stage 3 paralysis and complete absence of clasping reflex in both hindlimb (O). **(P)** Comparison of paralysis stages over time in TauP301S (n=4) and cDKO; TauP301S (n=5) mice. Two-way Anova followed by Sidak’s test, * p≤0.05, **p≤0.01. **(Q)** Percentage of mice at stage 3 over time in cDKO;Tau WT (n=6), TauP301S Tg mice (n=32), and cDKO; TauP301S (n=14). Mantel-Cox test ****p≤0.0001. Inset shows representative image of hunched spine in the hindlimb region of cDKO; TauP301S at 350 days compared to aged-matched TauP301S.

We next wondered whether the pro-survival activity of Numb in tauopathies extends beyond RGCs to other neuronal cell types. To address this question, we capitalized on the fact that the Islet1-Cre driver used to generate the cDKO mouse line is also active in developing spinal motoneurons from E10.5(*55–57*) and in mature motoneurons of the medial lateral motor column (LMC_M_) of the spinal cord (*58*), allowing for inactivation of Numb in these cells. Interestingly, TauP301S mice were also previously reported to exhibit progressive spinal motoneuron degeneration and clasping of the hindlimb, which eventually progresses to complete paralysis and inability to feed (*59–64*). This provided an opportunity to test whether loss of Numb would accelerate motoneuron degeneration and the appearance of motor phenotypes in TauP301S mice.

We first sectioned spinal cords in the L3 and L4 lumbar region and counted the number of motoneurons in the LMC_M_ by staining for Choline acetyl-transferase (ChAT), a pan-motoneuron marker. In cDKO; TauWT mice, we found similar numbers of motoneurons at 600 days as in WT mice at 350 days (Fig. S4 D, E-G), indicating that, unlike in the retina, Numb and Nbl are not essential for long-term survival of motoneurons in the context of WT Tau expression. In TauP301S mice, we found a decrease in the number of ChAT+ motoneurons compared to WT at 325-350 days, as previously reported(*60*), but this decrease was significantly more important in cDKO; TauP301S mice at both the L3 and L4 level (Fig. 4F–L). Consistently, histological analysis of the spinal cord showed a reduction of large soma neurons in the LMC_M_ of cDKO; TauP301S compared to TauP301S (Fig. S4A-C), indicating that the reduced number of ChAT+ cells reflect motoneuron loss rather than downregulation of ChAT. These results indicate that Numb/Nbl normally function to delay spinal motoneuron degeneration in the context of tauopathy.

To study motor function, we tested the hindlimb clasping response at different ages as an indicator for the severity of paralysis (*65, 66*). We defined three stages of clasping depending on the distance observed between the hindlimbs when raising the mouse by the tail (Fig. 4M), as previously described (*67*). We followed mice in the TauP301S and cDKO; TauP301S group over time and observed that stages 2 and 3 are reached on average significantly earlier in cDKO; TauP301S mice (Fig. 4N–P), and 100% of cDKO; TauP301S animals displayed a stage 3 clasping response by 390 days, whereas it took 501 days to reach this stage in 100% of TauP301S mice (Fig. 4Q). The cDKO; TauP301S animals also presented a hunched spine in the hindlimb region (Fig. 4Q), and failed to grab the cage in an upside-down position at a younger age than TauP301S mice (Videos S1 and S2), consistent with the precocious development of the clasping phenotype. As expected, based our observation that motoneurons do not degenerate in cDKO; TauWT animals (Fig. S4 D-G), we observed no paralysis in these animals (Fig. 4Q). Together with our observations in the retina, these results indicate that Numb promotes neuronal survival in the context of tauopathy in multiple neuronal cell types.

### Numb negatively regulates intracellular Tau levels in an isoform-specific manner

As inactivating Numb results in increased Tau levels and neurodegeneration in both WT and tauopathy mice, we postulated that elevating Numb might conversely reduce Tau levels and promote neuronal survival. Given the known role of Numb as an adaptor protein, we first asked whether it could physically interact with Tau. We found that each of the four isoforms of Numb co-immunoprecipitate with Flag-tagged Tau when expressed in HEK293T cells (Fig. 5A, B), consistent with physical interaction. To determine whether Numb regulates intracellular Tau levels, we first co-expressed each isoform of Numb together with Tau∷GFP in HEK293T cells and analyzed the levels of Tau∷GFP by Western blot 72 hours after transfection. While Numb-65, Numb-66 and Numb-71 had no effect, Numb-72 appeared to reduce the levels of Tau∷GFP (Fig. 5C). To provide a more quantitative measurement of this effect, we used a human medulloblastoma-derived cell line (DAOY) expressing dsRed together with Tau∷EGFP downstream of an internal ribosomal entry site (IRES)(*15*). This cell line allowed us to test the effects of Numb expression on Tau levels by reading out GFP fluorescence intensity, while using dsRed as a stable fluorescence signal for normalization (Fig. 5D). We transfected DAOY cells with constructs expressing Myc-tagged versions of each of the Numb isoforms or Myc alone as a control, and analyzed the relative levels of Tau∷EGFP and dsRed by flow cytometry 72 hours later. We found that expression of Numb-72 specifically increases the proportion of cells with low levels of Tau∷EGFP, whereas expression of Numb-65, Numb-66 and Numb-71 has no effect compared to Myc (Fig. 5E–J; S5A). These results indicate that elevating Numb-72 expression is sufficient to reduce intracellular Tau levels.

**Fig. 5.**
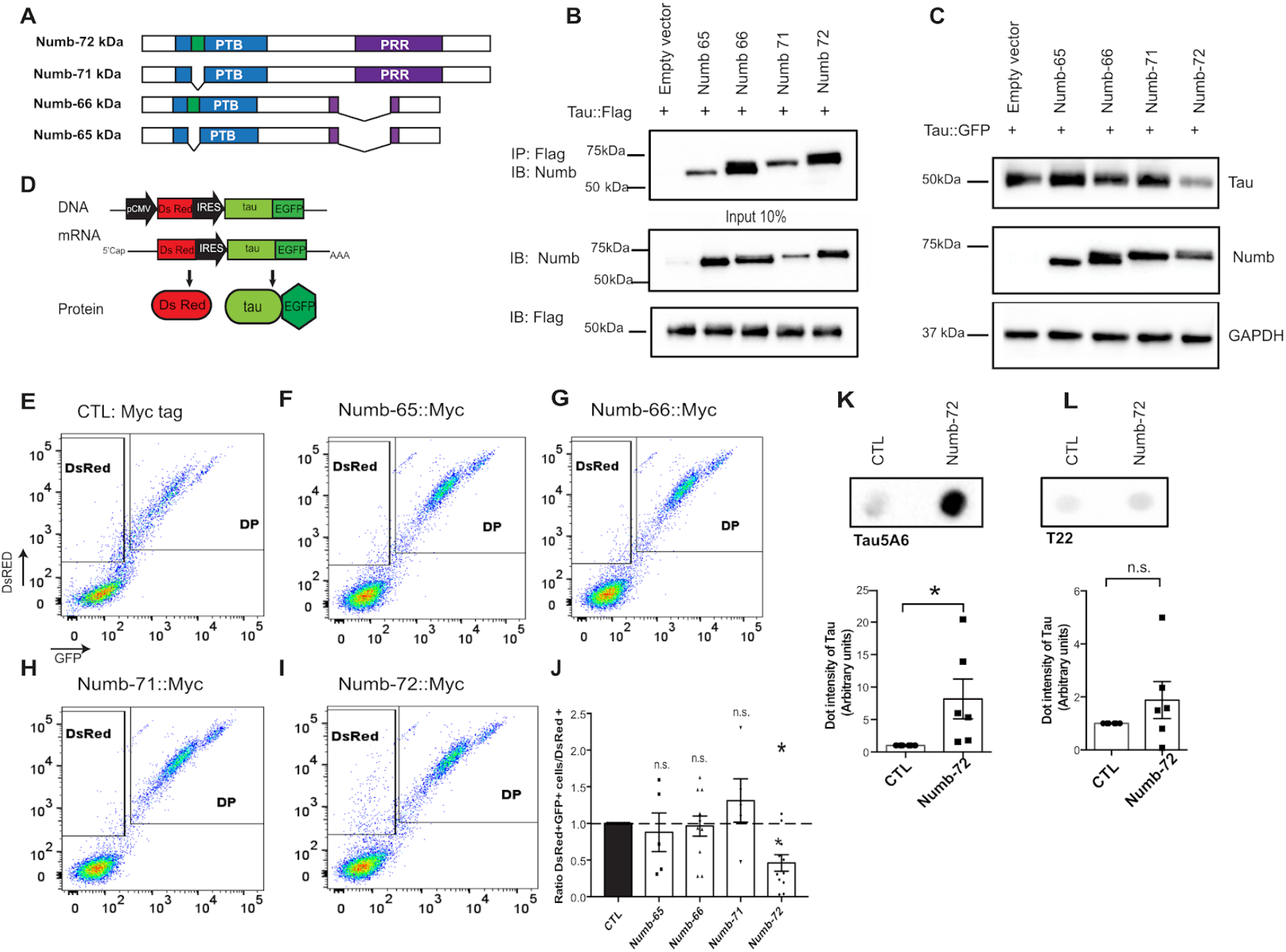
Numb-72 negatively regulates intracellular Tau levels. **(A)** Schematic representation of the four Numb isoforms, differing by the presence or absence of an insertion in the phosphotyrosine domain (PTB, green box) or the proline-rich region (PRR). **(B)** Coimmunoprecipitation (IP) of flag-tagged Tau (Tau∷Flag) with each isoform of Numb after co-expression in HEK293 cells. 10% input from protein extracts immunoblotted (IB) for Numb or Flag are shown as controls. Numb-specific bands migrate at ~72, 71, 66, and 65kDa. Tau∷Flag band migrates at ~50kDa. **(C)** Western blot analysis of Tau∷GFP (top blot) and Numb (middle blot) levels in HEK293 cells 72 hours after co-expression with the different Numb isoforms. GAPDH is used as loading control. **(D)** Schematic representation of the DNA insertion in the stable human-medulloblastoma-derived cell line (DAOY) and the resulting mRNA and protein expression in this cell line(*15*). **(E-I)** Flow cytometry analysis of the DsRed and GFP signal in the DAOY cells 3 days after transfection with constructs expressing each of the four different Numb isoforms or the Myc tag alone (control, CTL). The boxed area indicates the gating used to identify the DsRed+/GFP- population. DP: double-positive (DsRed+/GFP+) cells. **(J)** Average ratio of DsRed+/GFP+ cells over DsRed+ cells after expression of the Myc tag control (CTL) and the different Numb isoforms. The Myc tag value was normalized to 1 and used for comparison with all other conditions. Graph shows mean +/− SEM, *p ≤0.05; n.s.: not significant. One-way Anova followed by Dunett’s test. n=11 Myc, n=5 Numb-65, n=11 Numb-66, n=5 Numb-71 and n=11 Numb-72. N represents number of independent experiments. **(K, L)** Dot blot analysis of total Tau (Tau5A6 antibody, K) and oligomeric Tau (T22 antibody, L) levels detected in the culture medium of Tau-expressing HEK293T stable cell line 24 hours after transfection with either GFP (Control) or Numb-72-IRES-GFP (Numb-72). Graphs show dot intensity quantification of total Tau or oligomeric Tau signal in dot blot assays. Bars show mean ± SEM, n.s.: not significant; *p < 0.05. Student’s t test, n = 6 independent experiments.

To explore how Numb-72 regulates intracellular Tau levels, we first tested whether it could promote proteasome- or lysosome-mediated degradation of Tau (*68*). We transfected DAOY cells with constructs expressing Myc-tagged or Numb-72∷Myc, blocked the proteasome or lysosome 48 hours after transfection, and measured the levels of Tau∷EGFP and dsRed 24 hours later by flow cytometry. We predicted that if Numb-72 promotes Tau∷EGFP trafficking to the proteasome or lysosome, blocking these pathways would abrogate the effect of Numb-72. We observed that expression of Numb-72 decreases the number of cells with high levels of Tau∷EGFP, as observed in our previous experiments, but blocking the proteasome or lysosome has no effect on this activity (Fig. S5B), arguing against our hypothesis. Another pathway involved in the regulation of Tau levels is autophagy (*69*), and Numb was previously described to regulate this process (*70*). We therefore analyzed the ratio of autophagy markers LC3-I and LC3-II (*71*) in cDKO optic nerve extracts, but found no difference compared to Nbl KO controls, both in native (Fig. S5C, E) and stressed conditions after NMDA injection (Fig. S5D, F). Finally, we wondered whether Numb might promote the extracellular release of Tau, which is known to help balance intracellular levels in physiological and pathological conditions (*72, 73*). We transfected GFP or Numb-72 in a Tau-expressing stable cell line and measured the levels of total Tau and oligomeric Tau in the culture medium 24 hours later by DotBlot assays. Interestingly, we found that expression of Numb-72 leads to an increase in total Tau levels in the medium (Fig. 5K), but has no effect on the levels of oligomeric Tau released (Fig. 5L). These results suggest that Numb-72 promotes the extracellular release of native Tau, thereby leading to reduction of intracellular levels.

### Numb-72 prevents neurodegeneration in mouse models of tauopathy

The identification of Numb-72 as a negative regulator of intracellular Tau levels suggested that it might function as a neuroprotective factor in tauopathies. To test this idea, we first cultured primary RGCs from TauP301S transgenic mice and from the triple transgenic mouse model of Alzheimer’s disease (3xTg), which expresses the human mutant version of TauP301L, in addition to the Swedish APP mutant, and M146V Presenilin. RGCs were electroporated with constructs expressing GFP alone or Numb-72-IRES-GFP, cultured for two weeks and stained for NF165 to identify RGCs. While neurite length and number of branches were not changed in any condition (Fig. 6F, G, M, N), we observed a significant increase in the number of blebs in GFP-transfected TauP301S or 3xTg neurons (Fig. 6E, L), consistent with the idea that blebbing is an early sign of neurodegeneration in tauopathies. In contrast, the number of blebs in TauP301S and 3xTg neurons transfected with Numb-72 was not different from that of WT RGCs transfected with GFP (Fig. 6E, L), indicating that expression of Numb-72 prevents the appearance of neurodegenerative features in cultured neurons from two different mouse models of tauopathy.

**Fig. 6.**
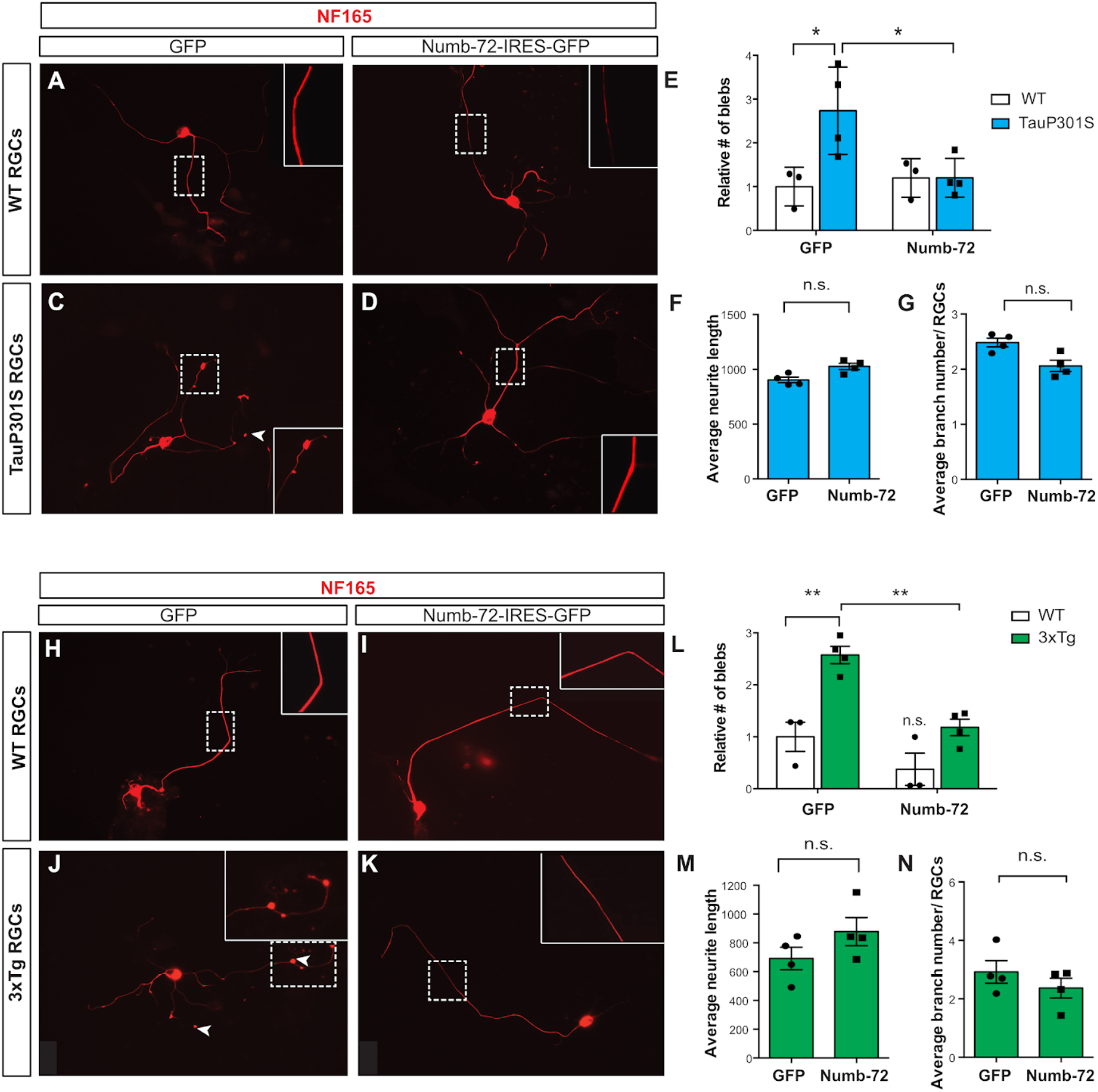
Numb-72 reduces blebbing in primary RGCs of tauopathy mouse models. **(A-D)** Immunostaining for neurofilament 165 (NF165) on primary retinal cell cultures from wildtype (WT) and Tau P301S mice 14 days after transfection with GFP or Numb-72-IRES-GFP. An increase in the number of blebs (arrowheads) is observed in TauP301S neurons compared to WT, but reversed by Numb-72 expression. Dotted boxes identify regions magnified in insets. **(E-G)** Quantification of the number of axonal blebs (E); neurites length (F) and branch number (G) in WT and TauP301S RGCs. Values in WT were normalized to 1. Bar graphs show mean ± SEM, ** p ≤0.001; n.s.: not significant. Two-way Anova followed by Sidak’s test (bleb counts), and Student’s t test (neurite length and branch numbers). n=3 independent cultures for WT (133 GFP-transfected cells and 125 Numb-72-transfected cells analyzed), and n=4 independent cultures for TauP301S (194 GFP-transfected and 121 Numb-72-transfected cells analyzed). **(H-K)** Immunostaining for neurofilament 165 (NF165) on primary retinal cell cultures from wildtype (WT) and 3xTg mice 14 days after transfection with GFP or Numb-72-IRES-GFP. An increase in the number of blebs (arrowheads) is observed in 3xTg neurons compared to WT, but reversed by Numb-72 expression. Dotted boxes identify regions magnified in insets. **(E-G)** Quantification of the number of axonal blebs (E); neurites length (F) and branch number (G) in WT and 3xTg RGCs. Values in WT were normalized to 1. Bar graphs show mean ± SEM, ** p ≤0.001; n.s.: not significant. Two-way Anova followed by Sidak’s test (bleb counts) and Student’s t test (neurite length and branch numbers). n=3 independent cultures for WT (128 GFP-transfected cells and 137 Numb-72-transfected cells analyzed), and n=4 independent cultures for TauP301S (151 GFP-transfected and 119 Numb-72-transfected cells analyzed).

Finally, we asked whether elevating Numb-72 levels in tauopathy neurons could prevent neurodegeneration *in vivo.* To deliver Numb-72 in RGCs, we generated AAV2 vectors expressing Numb-72 or GFP as control. As previously reported with AAV2, intravitreal injections preferentially infected RGCs (*47*) (Fig. 7A–C), and the Numb72-infected RGCs expressed Numb at levels well above baseline (Fig. 7D–F), validating this vector as an appropriate tool to elevate Numb-72 levels in RGCs *in vivo*. We injected AAV2-GFP or AAV2-Numb-72 in the vitreous of 5-month-old WT, TauP301S, and 3xTg mice. Seven weeks later, we induced a mild stress by intravitreal injection of NMDA in all mice (as in Fig. 3) and analyzed the number of RGCs three days later. In AAV2-GFP-infected retinas, we found that the number of RGCs is reduced by about 50% in TauP301S and 3xTg mice compared to WT (Fig. 7G, I, K, L, N, P), indicating that RGCs in these mouse models are more susceptible to stress-induced cell death, as previously demonstrated (*34, 36, 74, 75*). In AAV2-Numb-72- and AAV2-GFP-infected WT retinas, the number of RGCs was comparable, indicating that expression of Numb-72 does not prevent NMDA-mediated neuronal degeneration nor affect survival in WT neurons (Fig. 7G, H, K, L, M, P). Remarkably, however, there was no significant loss of RGCs in TauP301S or 3xTg mice infected with AAV2-Numb-72 compared to WT mice (Fig. 7H, J, K, M, O, P). These results indicate that viral vector-mediated delivery of Numb-72 prevents neuronal cell loss in two different mouse models of tauopathy *in vivo*.

**Fig. 7.**
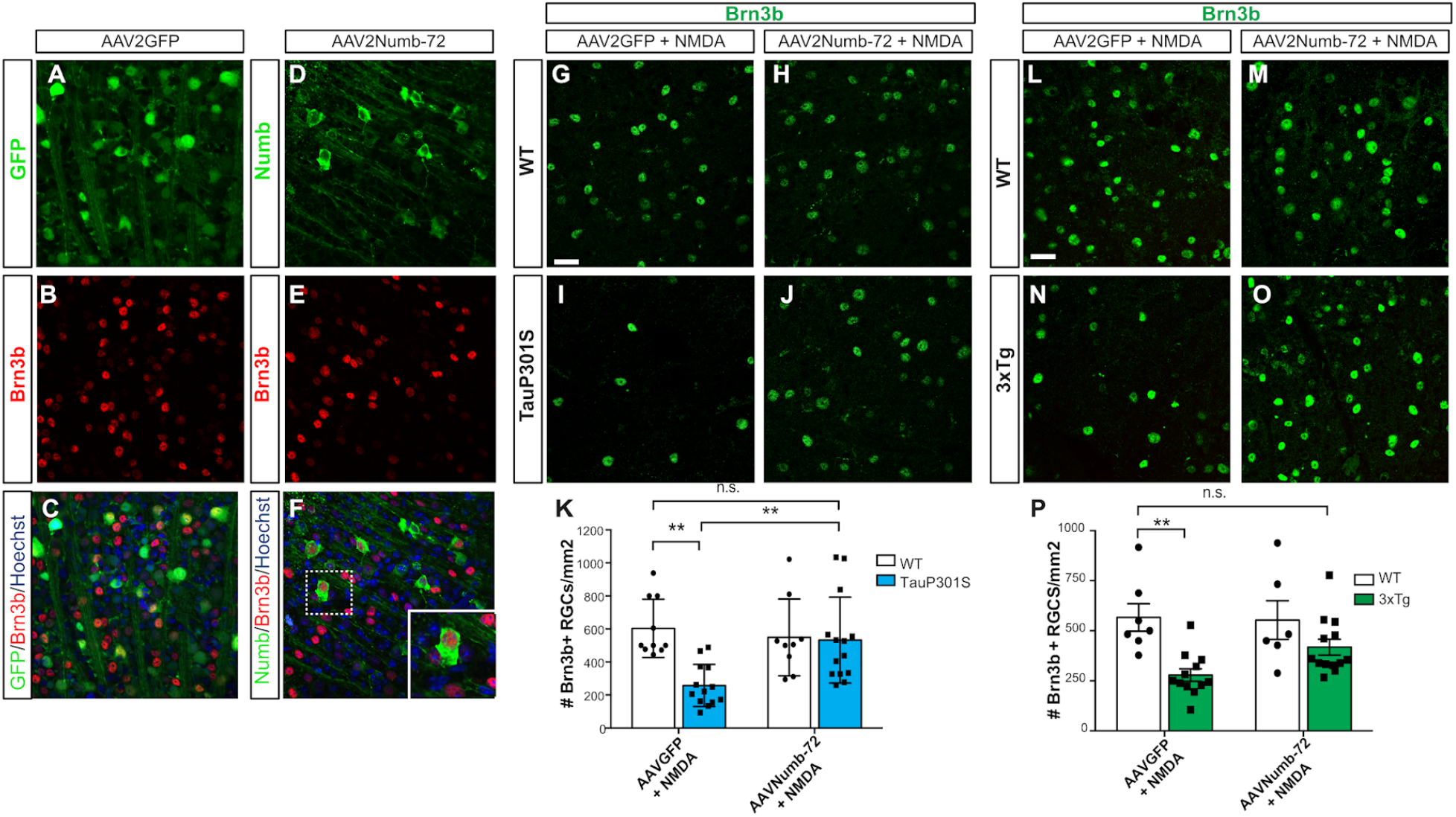
Numb-72 rescues RGC death in tauopathy mouse models *in vivo*. **(A-C)** Immunostaining for GFP (A) and Brn3b (B) on retinal flatmounts from WT mice, 7 weeks after intravitreal injections of AAV2-GFP in 5 months-old animals. Merged image together with Hoechst nuclear stain is shown in C. (D-F) Immunostaining for Numb (D) and Brn3b (E) of retinal flatmounts from WT mice, 7 weeks after intravitreal injections of AAV2Numb-72 in 5 months-old animals. Merged image together with Hoechst nuclear stain is shown in F. Inset shows high magnification view of a Numb/Brn3b double-positive cell. **(G-J)** Immunostaining for Brn3b on retina flatmounts from WT (G-H) and TauP301S mice (I-J), 7 weeks after intravitreal injections of AAV2-GFP or AAV2-Numb-72 in 5 months-old animals. Three days prior to sacrifice, all animals received an intravitreal injection of 10mM NMDA. Confocal images were taken in the ganglion cell layer. Scale bar = 30μm. **(K)** Quantification of the number of Brn3b+ RGC per mm^2^ in WT and TauP301S mice after AAV2-GFP + NMDA or AAV2-Numb-72 + NMDA injection. Bar graphs show mean ± SEM, n.s. = not significant, ** p ≤0.001; Two-way Anova followed by Turkey’s test. n= 10 WT+AAVGFP; n= 14 WT+AAVNumb-72; n= 7 TauP301S+AAVGFP; n= 14 TauP301S+AAVNumb-72 **(L-O)** Immunostaining for Brn3b on retina flatmounts from WT (L, M) and 3xTg-AD (N, O) mice, 7 weeks after intravitreal injections of AAV2-GFP or AAV2-Numb-72 in 5 months-old animals. Three days prior to sacrifice, all animals received an intravitreal injection of 10mM NMDA. Confocal images were taken in the ganglion cell layer. Scale bar = 30μm. **(P)** Quantification of the number of Brn3b+ RGC per mm^2^ in WT and 3xTg-AD after AAV2-GFP + NMDA or AAV2-Numb-72 + NMDA injection. Bar graphs show mean ± SEM, n.s. = not significant, ** p ≤0.001; Two-way Anova followed by Turkey’s test. n= 7 WT+AAVGFP; n= 12 WT+AAVNumb-72; n= 6 3xTg-AD+AAVGFP; n= 13 3xTg-AD+AAVNumb-72.

## DISCUSSION

Our findings indicate that Numb negatively regulates intracellular Tau levels, thereby preventing toxic accumulation of Tau oligomers and neurodegeneration. We also demonstrate that neuronal expression of the long isoform of Numb in two different mouse models of tauopathy prevents neurodegeneration, opening the door to the development of novel therapeutic approaches for tauopathies.

The role of Numb and Nbl has been extensively studied in the developing nervous system, where they control neural progenitor cell fate decisions in the neocortex (*41, 42, 76–78*), retina (*38, 79, 80*), and spinal cord (*23*). In contrast, their role in mature neurons is not as well-established. While Numb/Nbl inactivation disrupts axonal arborization in sensory neuron (*23*), axonal growth of hippocampal neurons (*24, 81*), and guidance of commissural neurons in the neural tube (*25*), these studies did not report an increase of neuronal cell death, which is in contrast with the present study. Importantly, previous studies did not evaluate neuronal cell loss in aged animals, whereas our data indicate that significant neuronal cell loss is not detected in the retina before 8 months in Numb/Nbl cDKO. Similarly, we previously reported that inactivation of Numb/Nbl in mouse rod photoreceptors leads to late-onset degeneration (*37*). In spinal motoneurons, however, we find here that inactivation of both Numb and Nbl has no effect on survival, even in two-year old animals, suggesting cell-specific functions under normal physiological conditions. In pathological contexts, however, such as in the TauP301S mice, inactivation of Numb/Nbl accelerates degeneration of both motoneurons and RGCs, suggesting a crucial pro-survival role when Tau levels are pathologically elevated. Loss of Numb/Nbl also accelerates the time of onset of motor deficits observed in TauP301S mice, consistent with the observed motoneuron loss. As the late onset of neurological phenotypes is often cited as a limitation for using Tau P301S mice as disease models, our findings suggest that Numb/Nbl cDKO; TauP301S mice might represent a more convenient model for some studies.

Our findings indicate that Numb can compensate for the loss of Nbl in RGCs, as Nbl inactivation alone is not sufficient to cause RGC death, even in old mice. Conversely, it remains unknown whether Nbl can compensate for the loss of Numb, as all mouse genetics experiments reported here were carried out in a Nbl KO background for simplicity. In several other contexts, however, mutual functional redundancy between Numb and Nbl was observed (*41, 42, 82*), including in the retina where Nbl was found to compensate for the loss of Numb in photoreceptor survival (*37*). Based on these observations, we speculate that Nbl could also compensate for the loss of Numb in RGCs, but addressing this question definitively will require further investigation.

Before neuronal cell loss, the first sign of degeneration detected in cDKO neurons is axonal blebbing, both *in vivo* and in culture, which is similar to what is observed in various neurodegeneration models (*45, 46*). Because inactivation of Numb in RGCs leads to elevation of Tau monomer and oligomer levels, we postulate that this elevation of Tau disrupts axonal transport, which eventually leads to cell death (Fig. 8A), consistent with previous findings showing a role for Tau in axonal transport (*83–85*). Our findings that overexpression of Tau in WT RGCs induces blebbing combined with results showing that knocking down Tau in cDKO RGCs rescues NMDA-mediated cell death also support a model in which elevation of Tau in cDKO RGCs is the cause of axonal blebbing and neuronal cell death.

**Fig. 8.**
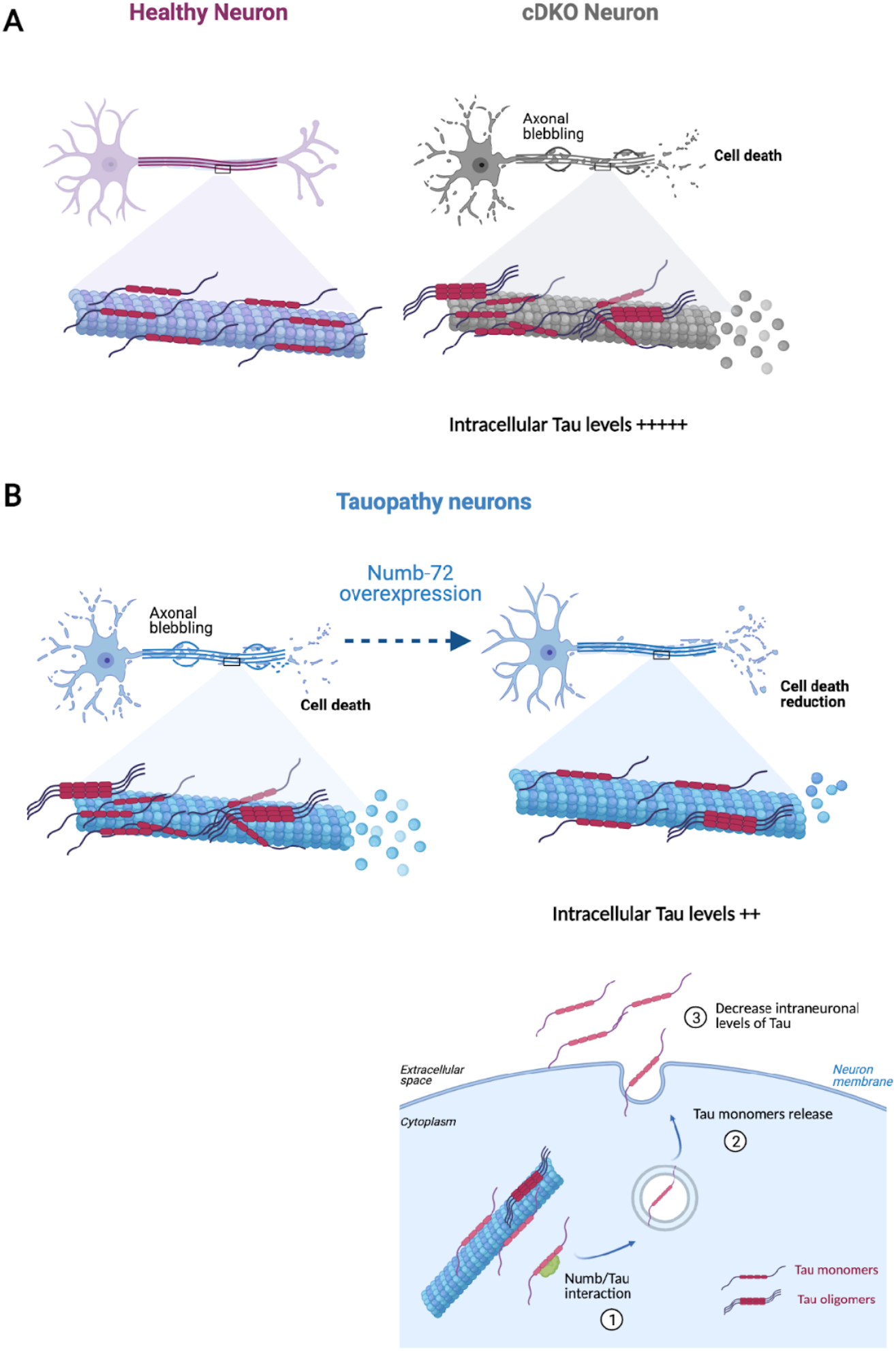
Model. **(A)** In healthy neurons (left), Numb/Nbl function to maintain Tau homeostasis and promote neuronal survival. In absence of Numb/Nbl (cDKO, right), intracellular Tau levels rise, leading to formation of Tau oligomers and axonal blebs, followed by cell death. (**B)** In tauopathy neurons, Tau oligomerize, which lead to axonal blebbing and neurodegeneration (left). Upon expression of Numb-72 in tauopathy neurons, however, the native form of Tau is released in the extracellular space, leading to reduced oligomeric Tau, fewer axonal blebs, and rescued neuronal survival (right).

Tau oligomers are the precursors of neurofibrillary tangles and are thought to play a crucial role in the onset and propagation of tauopathies (*86–89*). Our data show that loss of Numb leads to accumulation of both monomeric and oligomeric forms of Tau in the optic nerve, identifying a crucial role for Numb in the regulation of intraneuronal levels of Tau. While elucidating the precise molecular mechanisms by which Numb controls Tau levels will require further investigation, our findings indicate that Numb-72 promotes the release of native Tau monomers into the extracellular space. Several studies have shown that Tau is not only an intraneuronal protein, but can also be released in the extracellular space via exosomes and ectosomes (*72, 73, 90, 91*). Although the release of Tau oligomers was reported to contribute to spreading of the pathology, physiological Tau can also be secreted by neurons through exosomes (*92–94*), which is thought to serve as a clearance mechanism to prevent pathological Tau formation(*73*). Consistently, a recent report showed that the transcription factor EB (TFEB), a key regulator of lysosomal biogenesis, is essential for the exocytosis of specific Tau species (*95*). Inactivation of TFEB reduces the levels of Tau in the interstitial fluid in TauP301S mice and increases intracellular Tau levels, translating into enhanced pathology and spreading. By promoting the release of monomeric Tau in the extracellular space, we postulate that Numb-72 expression reduces the pool of Tau available in the cell to form oligomers, contributing to prevent cell death in tauopathies (Fig. 8B). Consistent with this idea, expression of Numb-72 in TauP301S RGCs, which have elevated oligomer levels due to Tau hyperphosphorylation, is sufficient to reduce the number of blebs in culture and fully rescue survival in vivo to the same level as WT RGCs. Expression of Numb-72 also decreases blebbing and neuronal cell loss in the 3xTg Alzheimer’s mouse model, albeit to a lesser extent. This observation is not surprising given that the 3xTg mouse also expresses mutated version of APP and Presenilin. But since Numb was found to interact with the cytoplasmic tail of APP and expression of Numb isoforms containing a PTB insertion (Numb-72 and Numb-66) reduces APP levels in SH-SY5Y cell line (*26, 96*), these results suggest that Numb-72 expression in 3xTg mice might have dual effect, lowering both Tau and APP levels.

While we cannot exclude that expression of Numb-72 in neurons might produce undesirable side effects, we have not observed any signs of toxicity when expressing Numb-72 in WT neurons. Nonetheless, it will be interesting in the future to carry out structure-function experiments to identify the minimal sequence of Numb-72 required to reduce Tau levels, as this may help generate a synthetic peptide that could be more suitable for clinical translation. Given that the only difference between Numb-72 and the other Numb isoforms lies in sequence insertions in the PRR and PTB domains, it is likely that these regions are involved. Future work will also help determine whether expression of Numb-72 can reduce Tau levels and prevent degeneration of different types of brain neurons, but our findings that inactivation of Numb accelerates neuronal degeneration in both the retina and spinal cord in mouse models of tauopathy suggests widespread pro-survival activity across the CNS.

## MATERIAL AND METHODS

### Animals

All animal work was carried out in accordance with the Canadian Council on Animal Care guidelines. The Islet1-Cre (*39*), *numb* floxed *(40) and numblike* null (*43*) mouse lines were used to generate Numb/Nbl cDKO. Female triple transgenic (3xTg) homozygote mice harbouring the APPSWE, PS1M146V and TauP301L transgenes and associated controls (female wild-type B6J129SF2J) were obtained from The Jackson Laboratory. Transgenic mice (PS19) expressing human TauP301S (1N4R), and the associated controls (wild-type C57b6J) were obtained from The Jackson Laboratory. We generated cDKO; TauP301S by intercrossing. TauP301S allele is hemizygous in all lines. Transgenic animals are always crossed with non-transgenics to maintain the line.

### Histology, immunohistochemistry

Eyes were enucleated and fixed by immersion in freshly prepared 4% paraformaldehyde (PFA) in phosphate-buffered saline (PBS) solution overnight (O/N) at 4°C, cryoprotected in sucrose 20% (in water) overnight, and cryosectioned at 14μm. For Numb staining at postnatal day 0 (P0), eyes were fixed in trichloroacetic acid 10% (TCA) for 10 mins and washed with PBS/30mM Glycine before cryoprotection in sucrose 20%. For retina flatmounts, eyes were enucleated after euthanasia and fixed for 2 h in 4% PFA before PBS wash (3 x 10 minutes) and dissected to isolate the neural retina, which was flatmounted on microscope slides with RGC facing the coverslip. Sections and flatmounts were pre-incubated for 1h in blocking solution (1% BSA in 0.4% Triton) and incubated overnight at 4°C with primary antibodies. Primary antibodies and conditions used in this study are listed in Table S1.

### RGC cultures

Retinal tissues were dissected from postnatal day 8 (P8) mice, cut in small pieces and incubated in PBS containing 5 mg/ml of papain (Worthington, LS003124), 0.24 mg/ml of L-cysteine, 0.5 mmol/l of EDTA, and 10 U/ml of DNase ┃ for 2×3 minutes. The enzymatic reaction was neutralized in a solution of low-ovomucoid-BSA (Sigma A-4161), trypsin inhibitor in PBS, and 10U/ml of DNaseI. Tissues were mechanically dissociated by gentle pipetting and cells collected as a suspension. Procedures were conducted at room temperature in a laminar flow hood. After centrifugation at 1000rpm for 11 minutes, cells were re-suspended in RGC medium: Neurobasal (Gibco) supplemented with Sato solution (Apo-transferrin (Sigma T-1147), BSA (Sigma A-4161), progesterone (Sigma P7505), putrescine, sodium selenite (Sigma S5261)), B27 supplement (Invitrogen), Penicillin/Streptavidin, Sodium Pyruvate (Gibco), Glutamax, N-Acetyl-Cystein (Sigma A8199), T3 (Triiodo-Thyronine) (Sigma T6397), Insulin (Sigma I6634), BDNF (Human Recombinant, Peprotec) and CNTF (Rat Recombinant, Peprotec). Cells were then plated on glass coverslips at 50 000 cells/coverslip in a 24 wells plate and incubated at 37°C; 8% CO2 for 2 weeks. Half of the culture medium was changed every 3 days. For RGC transfection, cells were re-suspended in Amaxa buffer and supplemented with 2μg of DNA (Numb-72 was cloned into pCIG-IRES-GFP, GFP was empty vector and Tau∷GFP and Tau∷P301L were gifts from Dr. N. Leclerc, U. Montreal) after centrifugation. Then, cells were transferred in an electroporation cuvette and nucleofection was carried out according to manufacturer’s recommendation (Amaxa nucleofector, program #33, Lonza Technology). Glass coverslips were incubated the day before with poly-D-lysine (PDL 10μg/ml) for 1h and laminin O/N.

### Generation of Tau stable cell line and cell culture conditions

A stable cell line was generated from Flp-In T-Rex 293 cells (Invitrogen), according to the manufacturer recommendations. 2μg of the POG44 vector in combination with 0.5μg of the vector pCDNA5-FRT/TO-*Tau* wa*s* transfected into Flp-In T-Rex 293 cells using the Jetprime Reagent (VRW). Following Hygromycin (Thermo Fisher) selection at 200μg/mL, a unique clone was selected.

HEK293T and stable cell lines (Daoy and TRex) were cultured in a humidified incubator at 37°C with 5% CO2 and fed every other day with high glucose DMEM medium (Thermo Fisher Scientific) supplemented with 10% FBS, 2mM glutamax (Thermo Fisher Scientific), 1mM sodium pyruvate (Thermo Fisher Scientific) and and 1% penicillin-streptomycin (10,000U/ml Thermo Fisher).

TRex cells at 50% confluence were treated with 1μg/mL tetracycline (Millipore-Sigma) for 24 h to induce protein expression before transfection for the dot blot assays.

### Flow cytometry analysis

The Daoy stable cell line expressing mCherry-IRES-Tau∷GFP(*15*) was grown in DMEM media to 50% confluence and transfected using Lipofectamine 2000 (Thermo Fisher Scientific) with expression constructs encoding: Myc, Numb-65∷Myc, Numb-66∷Myc, Numb-71∷Myc, and Numb-72∷Myc. To generate these constructs, appropriate cDNAs were cloned into the pCS2 plasmid. Cells were collected after 3 days, dissociated with 0.25% Trypsin-EDTA, and re-suspended in PBS. Samples were processed on a BD LSR Fortessa (BD biosciences) flow cytometer and analysis was done using FlowJo software. For protein degradation assays, cells were treated with MG132 (a proteasome inhibitor, 25uM Tocris #1748) or Chloroquine diphosphate salt (a lysosome inhibitor, 25uM Sigma-Aldrich # C6628) 48h after transfection.

### AAV production

AAV vectors were produced as previously described using the cotransfection method and purified by iodixanol gradient ultracentrifugation (*97*). AAV vector stocks were tittered by quantitative PCR (qPCR) (*98*) using SYBR Green (Thermo Fischer Scientific).

### Intraocular injections

For DiI retrograde labeling, eyes were enucleated and fixed overnight in 4% PFA at 4°C. The optic nerve was cut to within a few millimeters of the eye, and a few lipophilic DiI crystals (Invitrogen D282) were stuffed at the tip of the optic nerve into the optic nerve head. The eyes were then placed into fresh 4% PFA at 37°C for 7 days. Retinas were then dissected, flatmounted in Mowiol and imaged on an LSM710 (Zeiss) confocal microscope.

Saline, NMDA (N-Methyl-D-aspartic acid 10μM Tocris Bioscience), siRNAs (Dharmacon/Horizon) and AAVs were injected in adult eyes according to a modified procedure previously described (*99*). The titers of the AAV serotype 2 used were: 5.5 10e13 viral genome (vg)/ml for GFP = 1.1 10e11 vg/retina and 1.63 10e14 vg/ml = 3.26 10e11 vg/retina lot1 and 8.3 10e13 vg/ml= 1.66 10e11 vg/retina lot 2 for Numb72. The procedures for construction and purification of AAVs and intraocular injections were described previously (*19*). The volume of the injection was 2 μl. While under isofluorane anesthesia, animals were injected intravitreally with AAV-Numb72 in one eye, while injection of either vehicle or AAV-GFP into the contralateral eye served as an injection control. The eyes were collected 7 weeks after injections for AAV and after 3 days for the siRNA experiments. All eyes were fixed and processed for immunostaining on flatmount as described above. Scramble siRNA and siTau were generated by Dharmacon/Horizon Discovery group product ON-TARGETplus Smart pool siRNA Reagents – Mouse *Mapt* resuspended in RNase free water at 50μM and used at 10μM.

### Protein extraction, immunoblotting, and immunoprecipitation

For Western blots and immunoprecipitation, HEK293T cells were transfected with the Jet Prime (Polyplus transfection) using the following constructs: Tau∷Flag, Tau∷GFP (gifts from Dr. N. Leclerc, U. Montreal), Numb-65, Numb-66, Numb-71 and Numb-72 cloned into a pCIG2-IRES-GFP plasmid.

Cells were harvested and lysed in NP-40 buffer (50 mm TRIS, pH 8.0, 150 mm NaCl, 1.0% NP-40 with Complete Protease Inhibitor Cocktail from Roche). For Western blots, 40μg of protein extract (*in vivo*) or 10μg (*in vitro)* were loaded on 10% acrylamide SDS-PAGE gels for separation by electrophoresis and then transferred onto PVDF membranes using a transblot (Millipore). The membranes were blocked with 5% milk in Tris-Buffered saline solution with Tween (TBST, Tris-HCL concentration, NaCl concentration, pH 8.0, 0.1% Tween). Immunoblotting with the primary antibody was performed at 4°C overnight in 0.5% dry milk in TBST Primary antibodies and conditions used in this study are listed in Table S1. The primary antibody was detected with an HRP-conjugated goat anti-rabbit or anti-mouse (1:10,000; Jackson Immunoresearch) in 0.5% dry milk in TBST. HRP activity on the membrane was visualized with the ECL kit (GE Life Science). For quantifications, we used the Image Lab software (BioRad) to measure band intensity and used GAPDH band intensity for normalization.

For immunoprecipitation, Dynabeads Magnetic Beads (Dynabeads Protein G, Invitrogen) were used according to manufacturer’s specifications. Briefly, 40 μl of beads were incubated with primary antibody for 1h at 4°C. One mg of cell lysate was incubated with the bead-antibody mixture in Iph Buffer (50 mm Tris ph 8.0, 150 mm Nacl, 5 mm EDTA, 0.1% NP-40) overnight at 4°C. The beads were separated using a magnet (MagnaBind, Pierce) and washed in Iph Buffer. The beads were then boiled in 2× Laemmli buffer at 95°C for 10 min and the supernatant was used for immunoblotting as described above. We used 10% of cell lysates for input.

For dot-blot experiments, we collected culture medium from the Tau-expressing stable cell line 24h after GFP or Numb-72 transfection. Culture medium was collected and centrifuged to eliminate cell debris, then 400 μl of medium was loaded on nitrocellulose membrane. For immunoblotting, the membranes were blocked with 5% milk in TBST and incubated with the primary antibody at 4°C overnight in 0.5% dry milk in TBST (Tau5A6, T22- see Table S1 for dilution conditions). The primary antibody was detected with an HRP-conjugated goat anti-rabbit or anti-mouse (1:10,000; Jackson Immunoresearch) in 0.5% dry milk in TBST. HRP activity on the membrane was visualized with the ECL kit (GE Life Science).

### RNA isolation and quantitative PCR

Eyes were dissociated and collected into Qiagen Buffer RLT plus, and RNeasy microkit (Qiagen, 74004) was used to isolate RNA according to the manufacturer’s protocol with an additional 2 min of vortex in Buffer RLT. Isolated RNA was reverse transcribed using Superscript VILO Master Mix (Thermo Fisher Scientific, 11755050). cDNA was amplified by quantitative PCR using SYBR Green Master mix (Thermo Fisher Scientific, A25742). The primers used for Tau are: forward primer. 5’.CGCCCCTAGTGGATGAGAGA.3’ and reverse primer 5’.GCTTCTTCGGCTGTAATTCCTT.3’ and for GAPDH: forward primer 5’.TGCAGTGGCAAAGTGGAGAT.3’ and reverse primer 5’.ACTGTGCCGTTGAATTTGCC.3’. All primers were validated using a standard curve dilution of cDNA before the experiments were conducted.

### Hindlimb paralysis evaluation

Mice from each group were sacrificed as soon as they presented total paralysis. Mice from groups TauP301S and cDKO; TauP301S were followed each day from day 200 onward by lifting the animals by the tail to evaluate hindlimb clasping reflex (Stage 1 to 3) to determine the stage of paralysis until they reached full paralysis (stage 3).

### Spinal cord sampling and embedding

Mice were anesthetized and transcardially perfused with saline followed by 4% PFA. Spinal cords were removed and cut into cervical, thoracic, lumbar, and sacral segments. For immunohistochemistry, spinal cord tissues were incubated in 20% sucrose in PBS O/N at 4°C, embedded in OCT/Sucrose (1/1, vol/vol) and then frozen in liquid nitrogen and stored at −80°C. ChAT immunohistochemistry (see Table 1) was carried out on coronal sections of the lumbar spinal cord (16 μm) at the age of 11 months old (325-350 days), as described above. Image acquisition and motoneuron number quantification was done on lumbar sections (see quantification details below).

### Quantification

RGCs survival was assessed by counting Brn3b-positive cells on retina flatmounts within four areas of 212μm x 212μm around the optic nerve head and averaging the number of cells/μm2 per retina. Single plane images were taken with a LSM 710 confocal microscope (Zeiss).

Cell type quantification in retinal sections was done by averaging the total number of positive cells for specific markers in a 250μm stretch of the central and peripheral retina on four different retinal sections per animal. Images were taken on an LSM710 confocal microscope (Zeiss).

Length of individual isolated RGC neurites was measured on Image J by drawing the neurite with the polyline function. Blebs were counted manually. Images were taken with a DM6000 epifluorescence microscope (Leica).

For motoneuron counting, ChAT-positive cells located only on the ventral part of spinal cord were counted by applying a FFT Bandpass filter on 16-bit images acquired on a LSM710 confocal microscope (Zeiss) to reduce background. Positive cells were selected by a manually defined fluorescence intensity threshold and automatically counted using the “Analyze Particles” plugin of ImageJ. Minimums of three sections were counted at each level of the lumbar spinal cord that was analyzed per animal. The resulting cell counts were then averaged to give a representative number of the remaining number of motoneurons in that animal.

## Statistical analysis

Data analysis and statistics were performed using Prism 6 by two-way analysis of variance (Sidak), one-way analysis of variance (ANOVA) followed by a Dunett or Tukey *post hoc* tests, or by a Student’s *t*-test, as indicated in the legends. For the paralysis assay we used Mantel-Cox test. All quantifications shown are mean ± s.e.m, unless otherwise indicated. n represents number of biological replicates.

## SUPPLEMENTARY MATERIALS

Fig. S1: Numb expression in retinal ganglion cells and generation of a conditional Numb/Nbl double knockout mouse line

Fig. S2. Age-related neurodegeneration in Numb/Nbl cDKO retinas

Fig.S3. Phospho-Tau expression analysis

Fig. S4. Accelerated spinal motoneuron degeneration in Numb/Nbl cDKO on a TauP301S background

Fig. S5. Numb-72 expression reduces intracellular Tau levels in a degradation-independent manner

Table 1: list of antibodies and condition of using

Video S1: Mouse TauP301S 350days

Video S2: Mouse cDKO;TauP301S 350 days (Stage3)

## ACKNOWLEDGMENTS

We thank the IRCM core facilities for technical assistance, in particular: Jessica Barthe for mouse colony management, Dominic Filion for help with microscopy, Eric Massicotte and Julie Lord for help with flow cytometry, and Simone Terrouz for hematoxylin/eosin colorations. We thank Nicolas Stifani and Benoit Boulan for help with Image J plugin used for cell quantifications, Nicole Leclerc and Alexandre Desjardins for the Tau∷Flag and Tau∷EGFP constructs, and Cristian A. Lasagna-Reeves and Huda Y. Zoghbi for the DAOY cells. We are also grateful to all members of the Cayouette lab, past and present, for ongoing suggestions on this work. This study was supported by grants from the Alzheimer Society of Canada and the Canadian Institutes of Health Research (FDN-159936) to M.C., and LabEx LIFESENSES (ANR-10-LABX-65) and IHU FOReSIGHT (ANR-18-IAHU-01) to D.D. M.L. was supported by an IRCM Foundation-Jean Coutu Postdoctoral Fellowship, and M.C. is an Emeritus Research Scholar from the Fonds de recherche du Québec Santé (FRQS) and holds the Gaëtane and Roland Pillenière Chair in Retina Biology from the IRCM Foundation.

## AUTHOR CONTRIBUTIONS

Conceptualization: M.L. and M.C; Experimentation and data analysis: M.L., S.H., C.J, K.S, T.B, and J.C. AAV virus production: M.D and D.D. Manuscript writing and editing: M.L. and M.C. Supervision and funding: M.C.

