## Supplemental figures and Table 1 for "Numb reduces Tau levels and prevents neurodegeneration in mouse models of tauopathy in an isoform-specific manner"

### SUPPLEMENTARY FIGURES AND LEGENDS

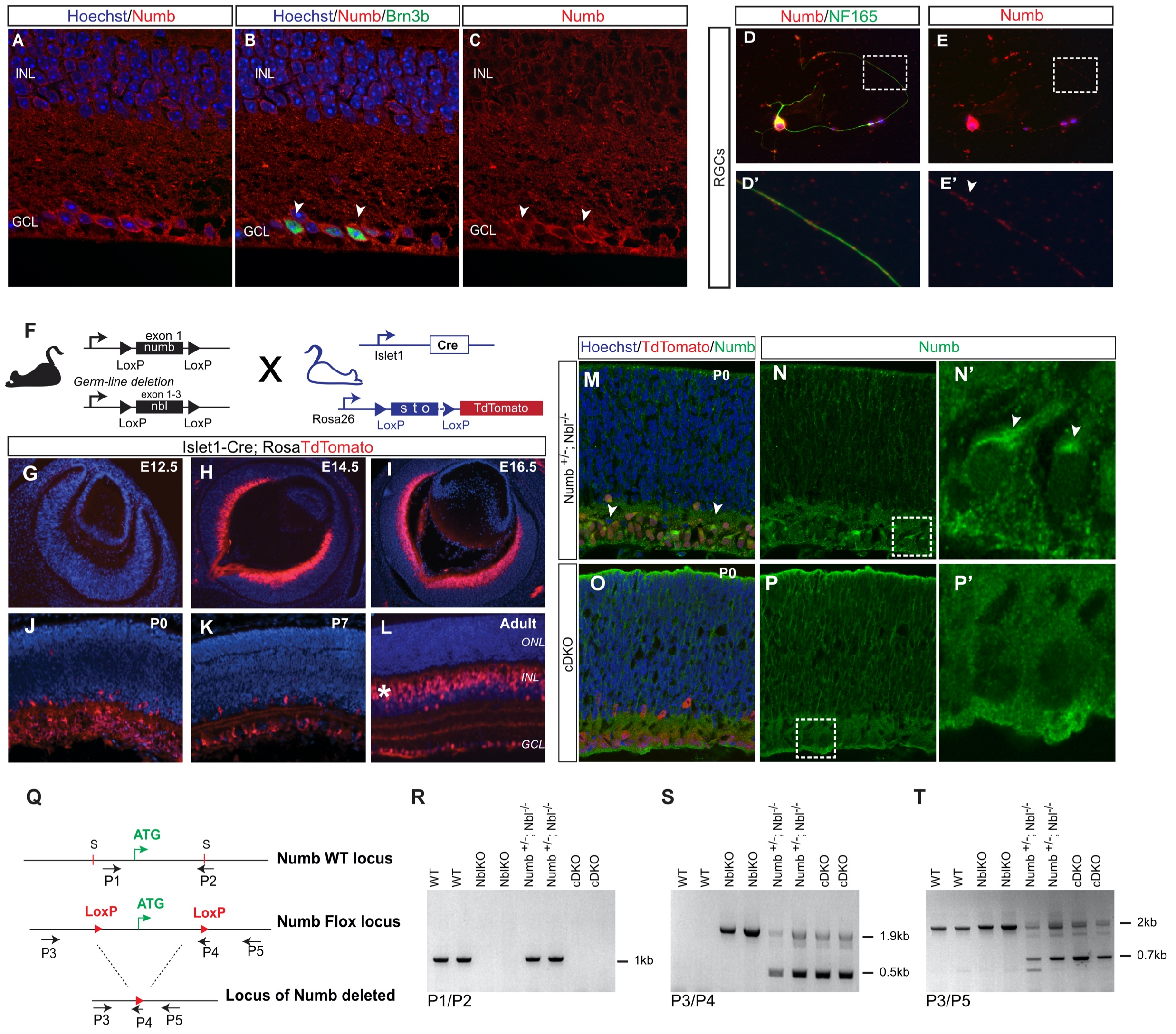


**Figure S1: Numb expression in retinal ganglion cells and generation of a conditional Numb/Nbl double knockout mouse line**

**(A-C)** Immunostaining for Numb (A) and Brn3b (B) on retinal sections from adult WT mice. Merged image in C includes Hoechst nuclear stain (blue). INL: inner nuclear layer. GCL: ganglion cell layer. **(D-E)** Immunostaining for NF165 (D) and Numb (E) in primary retinal cell cultures prepared from postnatal day 8 (P8) retina and cultured for 14 days. High magnification of the region identified by dotted line boxes are shown in D' and E'. Arrowheads indicates Numb expression in RGC axon. **(F)** Diagram illustrating the breeding scheme to generate Numb/Nbl cDKO in RGCs using the Cre/loxP system in a Numb-like (Nbl) null background and with Rosa-TdTomato mouse as a Cre reporter. **(G-L)** Cre activity in the retina of Islet1-Cre line using the Rosa-TdTomato reporter mouse at different stages (e12.5, e14.5, e16.5, P0, P7 and adult). Tdtomato is detected in developing RGCs from E14.5 (H, arrowheads), and bipolar cells in the adult retina (L, asterisk). **(M-P)** Immunostaining for Numb and TdTomato reporter on P0 retinal sections from $Numb^{-/+}$; $Nbl^{-/-}$ and cDKO animals. High magnification images of regions identified by dotted lines in N and P are shown in N' and P'. Arrowhead points to specific Numb signal in RGC soma. **(Q)** Diagram of the Numb WT and floxed allele. Numb deletion is generated by Cre-mediated excision of the first coding exon of the Numb gene. Start codon (ATG); loxP site (Red triangle) inserted at site (S) on WT locus. Different primers (P) used for genotyping are represented by arrows. **(R-T)** PCR genotyping on adult retina WT, Nbl KO, $Numb^{-/+}$; $Nbl^{-/-}$ and cDKO mice, using primer pairs P1 and P2 to detect WT allele (1kb band), P3 and P4 to detect flox allele (1.9kb band) and deletion when Ilset1Cre is active (0.5 kb band), and P3 and P5 to detect the Numb deletion allele (0.7 kb band).

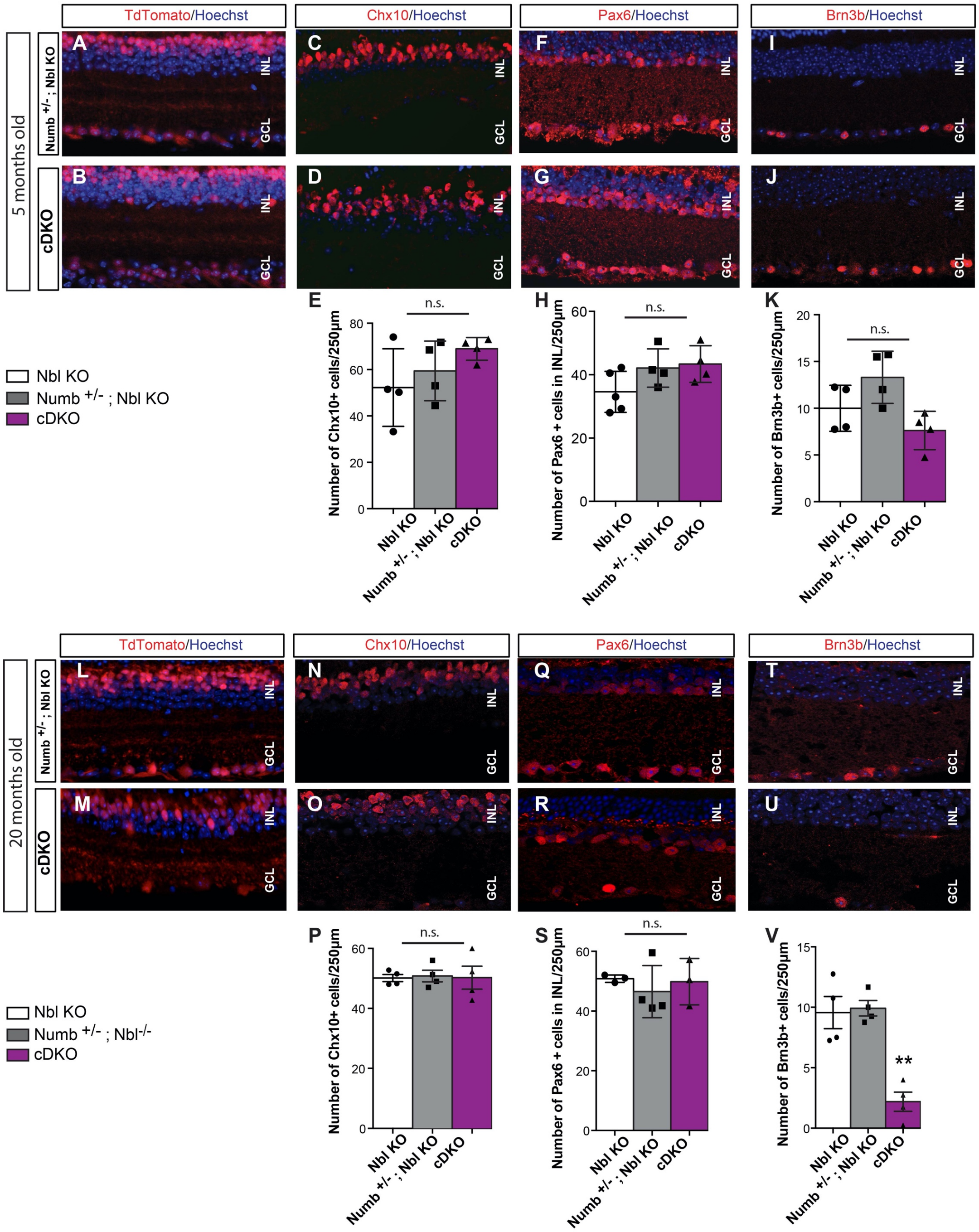


### Figure S2. Age-related neurodegeneration in Numb/Nbl cDKO retinas

**(A-H)** Immunostaining on retinal sections from 5-month-old Numb$^{-/+}$; Nbl KO and cDKO for TdTomato (A, B), CHX10 (C, D), Pax6 (F, G) and Brn3b (I, J). Nuclei are labelled with Hoechst (blue). **(I-K)** Quantification of the number of Chx10+ bipolar (E), Pax6+ amacrine (H) and Brn3b+ RGCs (K) in Nbl KO, Numb$^{-/+}$; Nbl KO, and cDKO. Mean ± SEM, n= 4 animals/genotype/time point. Anova test followed by Tukey's test. n.s: not significant. **(L-S)** Immunostaining on retinal sections from 20-month-old Numb$^{-/+}$; Nbl KO and cDKO for TdTomato (L, M), CHX10 (N, O), Pax6 (P, Q) and Brn3b (R, S). Nuclei are labelled with Hoechst (blue). **(T-V)** Quantification of the number of Chx10+ bipolar (T), Pax6+ amacrine (U) and Brn3b+ RGCs (V) in Nbl KO, Numb$^{-/+}$; Nbl KO, and cDKO. Mean ± SEM, n= 4 animals/genotype/time point. Anova test followed by Tukey's test n.s: not significant, **p≤0.01. INL: inner nuclear layer; GCL: ganglion cell layer.

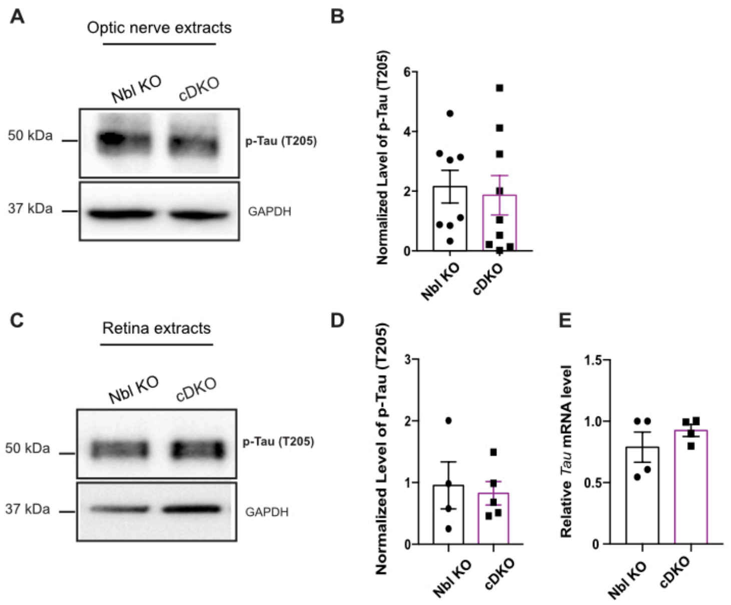


**Figure S3. Phospho-Tau expression analysis**

**(A-B)** Western blot analysis of phosphorylated Tau (T205) and GAPDH expression in optic nerve extracts prepared from 5-month-old Nbl KO and cDKO mice. **(B)** Quantification of the levels of Phospho-Tau in western blots, relative to GAPDH. Graph shows mean ± SEM, Student's t test, n.s. not significant, n= 8 NblKO and n= 9 cDKO. **(C-D)** Western blot analysis of phosphorylated Tau (T205) and GAPDH expression in retina extracts prepared from 5-month-old Nbl KO and cDKO mice. **(D)** Quantification of the levels of Phospho-Tau in western blots, relative to GAPDH. Graph shows mean ± SEM, Student's t test, n.s. not significant, n= 4 NblKO and n= 5 cDKO. **(E)** Relative level of Tau mRNA to GAPDH mRNA level in 5-month-old Nbl KO and cDKO retinal extracts measured by quantitative RT-PCR. Graph shows mean ± SEM, Student's t test, n.s. not significant, n= 4 NblKO and n= 4 cDKO.

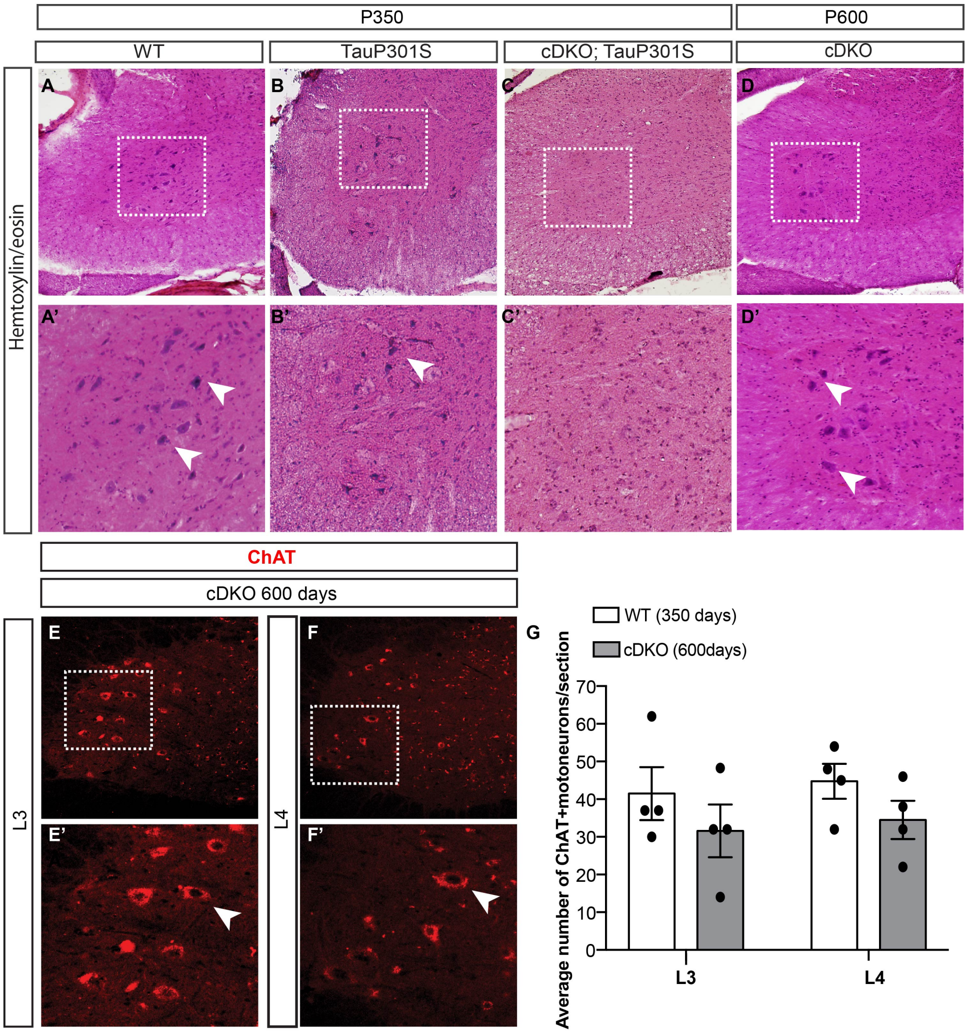


**Figure S4. Accelerated spinal motoneuron degeneration in Numb/Nbl cDKO on a TauP301S background**

**(A-C)** Hematoxyline-Eosin coloration on spinal cord sections of WT (A, A'), TauP301S (B, B') and cDKO; TauP301S (C, C') animals at 350 days. **(D-D')**. Hematoxyline-Eosin coloration on spinal cord section on cDKO animals at 600 days. Images in (A'-D') are high magnifications of the boxed regions in A-D. Arrowheads point to large soma neurons, a typical characteristic of motoneurons. **(E-F)**. Immunostaining for choline acetyltransferease (ChAT) on adult spinal cord sections in the lumbar region L3 (E, E') and L4 (F, F') on cDKO animals at 600 days. Images in E' and F' are high magnifications of the boxed regions in E and F. Arrowheads point to ChAT+ motoneurons. **(G)** Average number of ChAT+ motoneurons in lumbar region L3 and L4 in WT at 350 days and cDKO at 600 days. Mean ± SEM, n= 4 animals/genotype. Anova two-way test followed by Turkey's test, n.s. not significant.

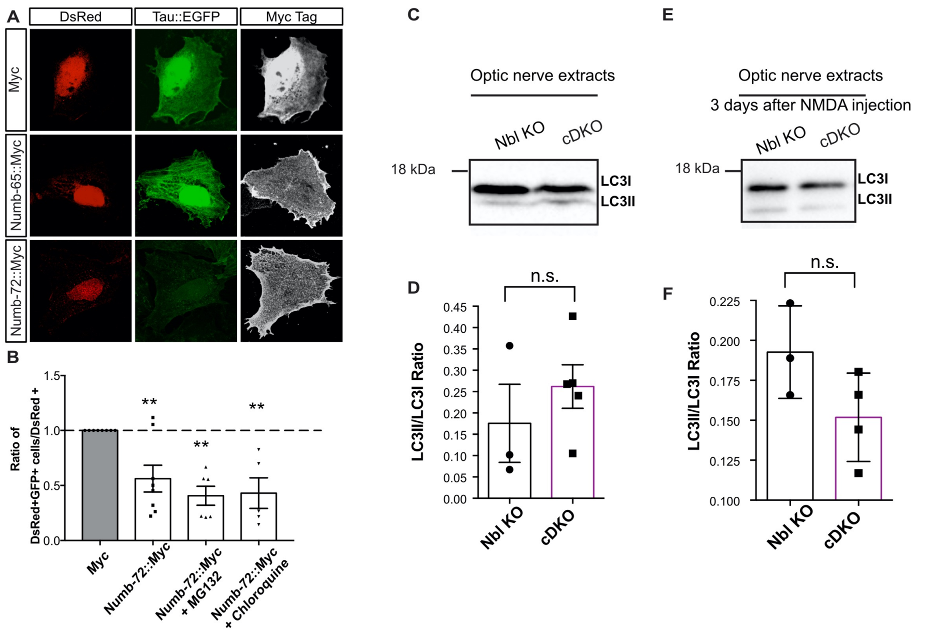


**Figure S5. Numb-72 expression reduces intracellular Tau levels in a degradation-independent manner**

**(A)** DsRed and Tau::EGF expression in DAOY cells transfected with constructs expressing Myc only, Numb-65::Myc, or Numb-72::Myc. The levels of Tau::EGFP are notably reduced after expression of Numb-72. **(B)** Average ratio of DsRed+/GFP+ cells over DsRed+ DAOY cells obtained by flow cytometry 3 days after transfection of the Myc tag alone or Numb-72::Myc. In the last and 24h, the culture medium was supplemented with DMSO alone or with MG132 (a proteasome inhibitor, 25uM) or Chloroquine (a lysosome inhibitor, 25uM) diluted in DMSO. The Myc tag value was normalized to 1 and used for comparison with all other conditions. Mean +/- SEM, **p ≤0.01; Anova one way test followed by Dunett's test, n= 8 Myc, 8 Numb-72::Myc, 6 Numb-72::Myc + MG132 and 6 Numb-72::Myc + Chloroquine. **(C, D)** Western blot analysis of LC3 and GAPDH expression in optic nerve extracts prepared from 5-month-old Nbl KO and cDKO mice without NMDA injection (C) or 3 days after NMDA injection (D). **(E, F)** Corresponding quantification of the ratio LC3II/LC3I in western blots for experiments shown in C and D. Graph shows mean ± SEM, Student's t test, n.s. not significant, dots on the graph represent independent biological replicates from different mice.

| Name | Cat# | Host | Source | Fixation | Dilution | Use |
|---|---|---|---|---|---|---|
| PAX6 | AB2237 | rabbit | Millipore-Sigma | 4%PFA, overnight @ 4C | 1 in 400 | IF |
| CHX10 | X1180P | sheep | Exalpha Biologicals | 4%PFA, overnight @ 4C | 1 in 400 | IF |
| BRN3B | SC-6026 | goat | Santa-Cruz Biotechnolog | 4%PFA, overnight @ 4C or 2h RT for flatmount | 1 in 500 | IF |
| Numb | ab14140 | Rabbit | Abcam | 4%PFA, overnight @ 4C or TCA 10%, 10min RT or 2h RT for flatmount | 1 in 200 (IF) and 1 in 3000(WB) | IF and WB |
| GFP | A11122 | rabbit | Thermo Fisher | 4%PFA, 10min RT (RGCs culture) | 1 in 1000 | IF |
| Neurofilamnt 165kDA (2H3) | AB 2314897 | Mouse | DSHB | 4%PFA, 10min RT (RGCs culture) | 1 in 200 | IF |
| Choline Acetyltransferase (ChAT) | AB 528122 | Mouse | DSHB | Perfusion 4%PFA + postfixation 4%PFA overnight @ 4C | 1 in 1000 | IF |
| MYC | SC-40 | mouse | Santa-Cruz Biotechnology | 4%PFA, 10min RT (Daoy cells) | 1 in 1000 | IF |
| Tau(K9JA) | A0024 | Rabbit | DAKO | 4%PFA, 10min RT (RGCs culture) | 1 in 5000 | IF |
| Tau (5A6) | AB_528487 | Mouse | DSHB | / | 1 in 1000 | WB and Dot Blot |
| Tau (T22) | ABN454 | Rabbit | Millipore Sigma | / | 1 in 1000 | WB and Dot Blot |
| Tau (T205) | ab254410 | Rabbit | Abcam | / | 1 in 1000 | WB |
| LC3b | 2775S | rabbit | Cell Signaling Technology | / | 1 in 1000 | WB |
| FLAG | 2368P | rabbit | Cell Signaling Technology | / | 1 in 1000 (WB), 1 in 50 (IP) | WB, IP |
| Glyceraldehyde-3-phosphate dehydrogenas | MAB374 | Mouse | Millipore Sigma | / | 1 in 5000 | WB |
| Acetylated tubulin | T6793 | Mouse | Sigma-Aldrich | / | 1 in 1000 | WB |

RT= Room Temperature
IF = Immunofluorescence
WB= Western Blot
IP = Immunoprecipitation
TCA= Trichloridacetic Acid
PFA= Paraformaldehyde
